# The selective autophagy receptor AtNBR1 associates with lipid droplets during seed germination and contributes to seed longevity

**DOI:** 10.64898/2026.11.23.768505

**Authors:** Isabelle Bouchez-Mahiout, Carine Deruyffelaere, Boris Collet, Marine Froissard, Loïc Rajjou, Anne Marmagne, Céline Masclaux-Daubresse, Sabine D’Andrea

## Abstract

Lipid droplets, once viewed primarily as lipid storage organelles, are increasingly recognized as key regulators of cellular homeostasis, which is particularly critical during seed imbibition and germination. Here, we investigated the role of the selective autophagy receptor AtNBR1 in lipid droplet-associated processes during Arabidopsis seed germination. AtNBR1 and ATG8 proteins were specifically and transiently recruited to lipid droplets during germination. Confocal imaging revealed their predominant colocalization in condensate-like puncta adjacent to lipid droplets and, less frequently, in autophagosome-like structures surrounding them. Mobile AtNBR1-positive lipid droplet remnants were also observed within the vacuole, particularly under treatments that simultaneously stimulate autophagy and limit vacuolar degradation. Besides, AtNBR1 degradation was modestly reduced in an *atg2* mutant, which is consistent with a minor NBR1-associated macrolipophagy-like pathway. Moreover, AtNBR1 does not appear to mediate a major route of lipid droplet degradation, as loss of AtNBR1 did not significantly affect bulk lipid turnover or the degradation of ubiquitinated and non-ubiquitinated oleosins, the major seed lipid droplet proteins, arguing against oleosins being primary AtNBR1 cargoes. Instead, our data support a broader role for AtNBR1 in seed proteostasis. AtNBR1 interactome comprises multiple protein quality control components, including the post-proteasomal exopeptidase TPPII. AtNBR1 also colocalized with CDC48A and PUX10, components of the lipid droplet-associated degradation pathway, and PUX10 contributed to AtNBR1 turnover. Importantly, loss of AtNBR1 compromised seed longevity in an accelerated aging assay. Together, our findings suggest a role for AtNBR1 in protein quality control at lipid droplets and support its positive contribution to seed longevity.

## Introduction

Lipid droplets (LDs) are evolutionarily conserved organelles that dynamically store and mobilize neutral lipids in response to metabolic and developmental cues, thereby providing substrates for energy production, membrane synthesis, and lipid signaling (Farese and Walther, 2025). Structurally, LDs consist of a core of neutral lipids, primarily triacylglycerols (TAGs), surrounded by a phospholipid monolayer that is associated with specific proteins that regulate their dynamics and functions. TAG mobilization from LDs is mediated by dedicated proteins through two main pathways: lipolysis, driven by cytosolic lipases, and lipophagy, mediated by autophagic pathways. Lipophagy encompasses distinct modes of LD degradation, including macrolipophagy, in which LDs are engulfed by autophagosomes that subsequently fuse with vacuoles/lysosome, and microlipophagy, in which LDs are directly engulfed by vacuoles/lysosome (Zienkiewicz and Zienkiewicz, 2020; Corbo and Chung, 2024; Zhao et al., 2025; Zhao et al., 2026).

In plants, cytosolic LDs are abundant in seeds, but also found in pollen and various vegetative organs (Bouchnak et al., 2023). LD degradation is best characterized during post-germinative seedling growth, where TAG mobilization is predominantly mediated by the LD-associated lipase SDP1 (Kelly et al., 2011; Zienkiewicz and Zienkiewicz, 2020). In addition to lipolysis, increasing evidence supports a role for lipophagy in plant LD turnover (Masclaux-Daubresse et al., 2020; Xu and Fan, 2022; Amari et al., 2024; Zhao et al., 2025). In rice cells, Kurusu et al. (2014) demonstrated an ATG7-dependent lipophagy process during anther development. Similarly, in Arabidopsis leaves, Barros et al. (2021) showed that, under carbon starvation, impairment of autophagy affects membrane lipid mobilization and prevents the formation of cytosolic LDs. Unlike in mammals, where macrolipophagy is the best-characterized form of lipophagy, current evidence indicates that microlipophagy-like processes may contribute to LD turnover in plants. In Arabidopsis leaves, Fan et al. (2019) described a lipophagy process resembling microlipophagy under carbon starvation. During seed germination, evidence for lipophagy remains limited to a proposed caleosin-dependent microlipophagy pathway (Poxleitner et al., 2006; Miklaszewska et al., 2023), leaving the contribution of selective autophagy to LD turnover unresolved.

Selective autophagy relies on cargo receptors that recognize specific substrates and connect them to the autophagy-related (ATG) machinery (Luo et al., 2021). Among these selective autophagy receptor (SARs), NBR1 (neighbor of BRCA1) is the most extensively characterized to date. Plant NBR1 is a functional hybrid of the mammalian autophagic adapters NBR1 and p62/SQSTM1 (Svenning et al., 2011). An increasing number of studies have revealed the role of NBR1 in plant responses to diverse stress conditions including heat, drought and nutrient starvation (Zhang and Chen, 2020) playing a central role in protein quality control (PQC). NBR1 is notably involved in the clearance of ubiquitinated protein aggregates, likely originating from denatured or damaged proteins generated by heat stress (Thirumalaikumar et al., 2020; Li et al., 2026). While NBR1 has been implicated in multiple degradation pathways in vegetative tissues, little is known about its function during seed germination. In Arabidopsis, it has been suggested that AtNBR1 may contribute to the modulation of abscisic acid signaling during imbibition and thereby influence seed germination (Tarnowski et al., 2020b). In addition, *Triticum aestivum* NBR1 has been implicated in enhancing drought tolerance during seed germination (Chen et al., 2022). Although plant NBR1 is best known as a selective autophagy receptor, recent findings suggest that its functions extend beyond autophagy and intersect with the ubiquitin–26S proteasome system (UPS). In rice, OsNBR1 interacts with proteasome components to promote the degradation of an auxin signaling regulator during root development (Sun et al., 2025), highlighting a role for NBR1 at the interface between autophagy and UPS-mediated proteostasis.

In germinating seeds, efficient LD mobilization requires not only TAG hydrolysis but also dynamic remodeling of the LD proteome, including the degradation of major structural proteins, such as oleosins, which contribute to LD stability. A specific pathway, termed LD-associated degradation (LDAD), mediates the dislocation of ubiquitinated oleosins from seed LDs. While K48-linked ubiquitination targets oleosins for degradation by the UPS, the fate and degradation mechanisms of oleosins carrying alternative ubiquitin linkages remain unclear (Deruyffelaere et al., 2018). Given the emerging evidence for lipophagy in plant LD turnover, this pathway may also contribute to the remodeling of the LD proteome.

Thus, despite the role of AtNBR1 in PQC and its potential involvement in seed germination, whether this selective autophagy receptor contributes to seed LD dynamics remains unknown. The recent identification of roles for mammalian NBR1 and p62/SQSTM1 in lipophagy further underscored the relevance of this question (Rasineni et al., 2017; Wang et al., 2017; Robichaud et al., 2021). Here, we investigated the role of AtNBR1 during Arabidopsis seed germination using complementary biochemical, imaging and proteomic approaches. We identify AtNBR1 as an LD-associated protein during early germination and show that its contribution to lipophagy is limited. Instead, our findings uncover an unexpected connection between AtNBR1, the LDAD machinery and PQC, suggesting that the turnover of LD-associated AtNBR1 is coordinated by the interplay between autophagic and LDAD/UPS pathways. Finally, we reveal a previously unrecognized role for AtNBR1 in maintaining seed longevity.

## Results

### AtNBR1 transiently accumulates in seeds after imbibition

To investigate the fate of AtNBR1 during seed development, we analyzed AtNBR1 levels in dry and germinating seeds by immunoblotting. Moist cold stratification, which releases seed dormancy in Arabidopsis, induced a progressive accumulation of AtNBR1 (Fig. 1A) which increased by 66% after 48 h of cold stratification (Fig. 1B). Consistently, analysis of *NBR1* expression in dry and 48 h-stratified seeds by reverse transcription-quantitative polymerase chain reaction (RT-qPCR) showed a significant increase in *NBR1* mRNA during seed stratification (Fig. 1C). Thus, under our experimental conditions, moist cold imbibition induces *NBR1* expression and AtNBR1 accumulation in seeds. Interestingly, ATG8 proteins (ATG8s), which interact with NBR1 and play a major role in the autophagy pathway, exhibited a dynamic similar to that of AtNBR1: their levels increased during cold stratification and subsequently decreased during germination (Fig. 1A). To determine whether AtNBR1 accumulation requires cold exposure, AtNBR1 content was analyzed in water-soaked seeds germinated at 25°C without prior cold stratification. After 24 h of germination under these conditions, AtNBR1 levels increased compared to dry seeds, although the difference was not statistically significant (Fig. 1B). These observations suggest that seed imbibition and the onset of germination promote AtNBR1 accumulation, while cold exposure may not be essential. When stratified seeds were exposed to light at 25°C to trigger germination, their AtNBR1 content decreased during germination (24 h) and post-germinative growth (48 h) (Fig. 1A). Taken together, these results indicate that AtNBR1 transiently accumulates in seeds at the initiation of germination.

**Figure 1.**
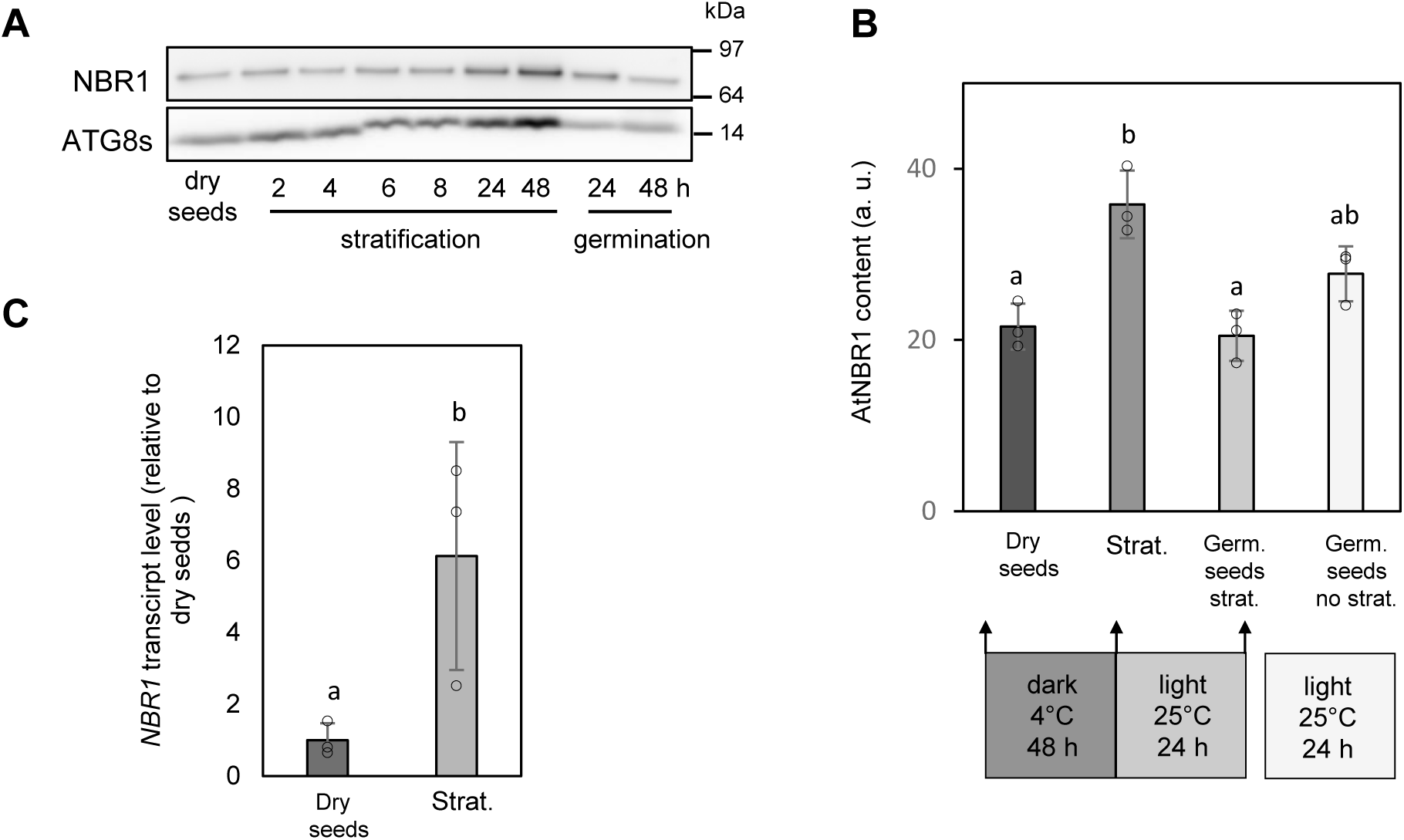
AtNBR1 transiently accumulates in Arabidopsis seeds at the initiation of germination. **(A)** AtNBR1 and ATG8s accumulate during seed stratification and are subsequently degraded during germination. Proteins were extracted from wild-type dry seeds, 2 to 48 h cold-stratified seeds, or 48 h-stratified seeds germinated for 24 and 48 h at 25°C. Proteins extracted from the same number of seeds were loaded on each lane and analyzed by immunoblotting with anti-NBR1 and anti-ATG8 antibodies. **(B)** Proteins from wild-type dry seeds, 48 h-stratified (Strat.) seeds, 48 h-stratified seeds germinated for 24 h (Germ. seeds strat.) or 24 h-germinated seeds without prior stratification (Germ. seeds no strat.) were extracted and analyzed as in (**A**). AtNBR1 content was measured using quantitative immunoblot (mean +/- SD from n = 3 samples of 100 seeds from independent plants). Significant difference from dry seeds was estimated using an exact one-tailed Mann-Whitney U Test (p=0.05). **(C)** AtNBR1 accumulation in stratified seeds is transcriptionally regulated. The expression of *NBR1* was analyzed by RT-qPCR in wild-type dry and 48 h-stratified seeds using specific primers NBR1-F and NBR1-R. The primer sequences are listed in Table S2. Normalization of *NBR1* transcript abundance was performed using *ASAR1* and *MAP2B* as reference genes (mean +/- SD from 3 samples of independent plants). Significant difference from dry seeds was estimated using an exact one-tailed Mann-Whitney U Test (p=0.05). Individual values are represented by open circles, and significant differences are indicated by different letters. **<u>Alt text</u>:** Figure 1: AtNBR1 transiently accumulates in Arabidopsis seeds at the initiation of germination; Figure 1A: AtNBR1 and ATG8s accumulate during seed stratification and are subsequently degraded during germination; Figure 1B: Moist cold imbibition induces AtNBR1 accumulation in seeds; Figure 1C: NBR1 transcripts accumulate in stratified seeds, showing that AtNBR1 accumulation is transcriptionally regulated.

### AtNBR1 and ATG8s mainly associate with LDs in dry and germinating seeds

To identify the intracellular localization of AtNBR1, we performed subcellular fractionation of dry, stratified, germinating and post-germinative seeds by density-gradient centrifugation, followed by immunoblot analysis of AtNBR1 content in the fractions (Fig. 2A). We also analyzed the distribution of several organelle markers in these fractions to identify the cellular compartments where AtNBR1 might be localized. In dry, stratified seeds and during early germination, AtNBR1 was predominantly found in the least dense fraction, which contains LDs as identified by OLE1, a specific LD marker. Interestingly, after 48 h of germination, the relative abundance of AtNBR1 in the LD-rich fraction decreased in favor of denser fractions, showing a distribution profile similar to that of PEX14, a peroxisomal marker, and VDAC1, a mitochondrial marker. In contrast, we observed a clear discrepancy between the localization of AtNBR1 and that of ARF1, the Golgi marker. These observations suggest that AtNBR1 is primarily associated with LDs in both dry and germinating seeds, but no longer during the post-germinative growth phase. To further explore the link between autophagy and LDs, we monitored the subcellular distribution of ATG8s. Fig. 2A shows that, like AtNBR1, ATG8s predominantly cofractionated with LDs in dry, stratified and germinating seeds, but not in post-germinative seedlings. Overall, these results suggest that AtNBR1 and ATG8s, two key players in selective autophagy, are transiently associated with LDs in dry and germinating seeds.

**Figure 2.**
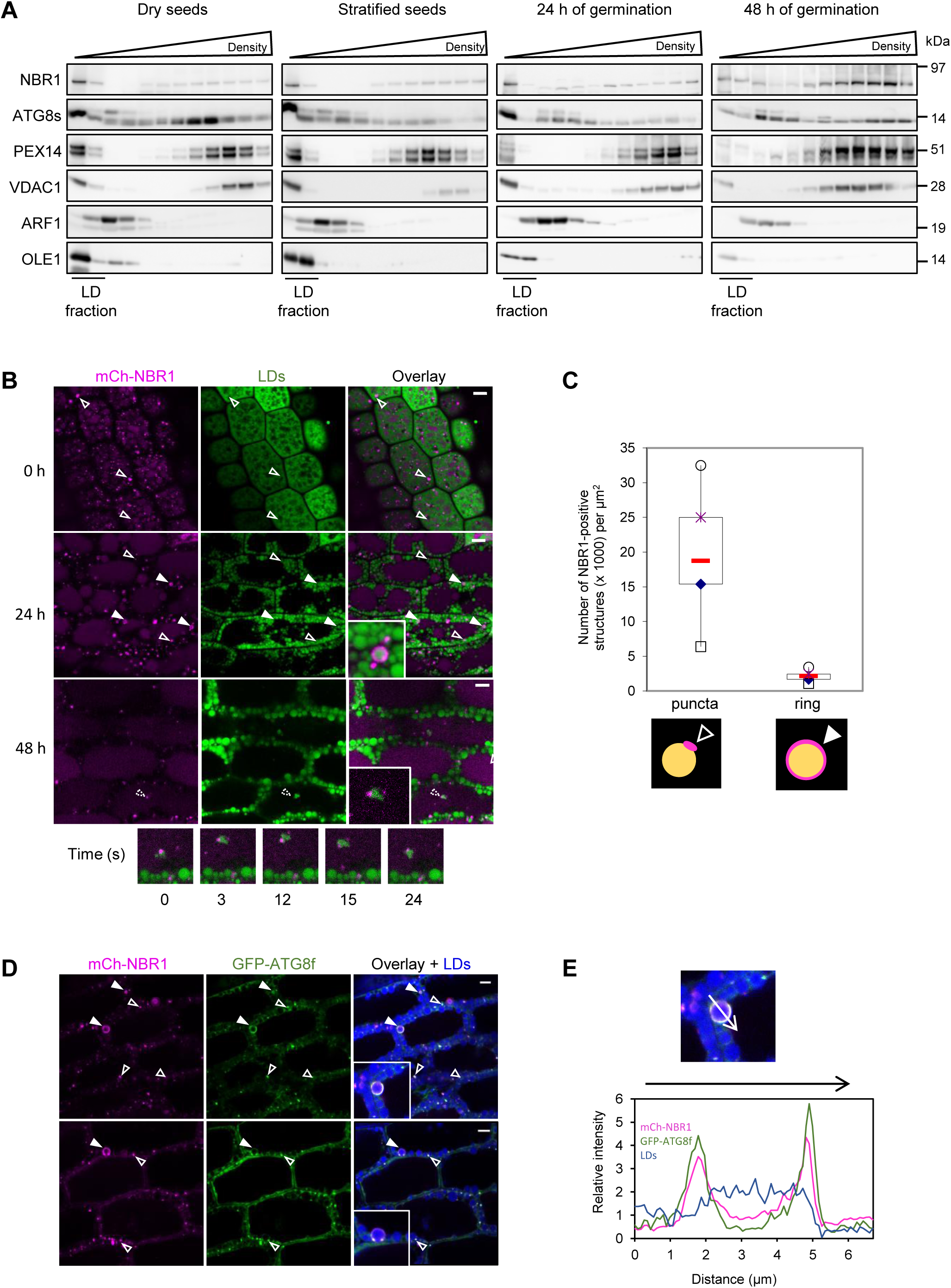
AtNBR1 and ATG8f associate with LDs via two distinct structures during seed germination. (**A**) AtNBR1 and ATG8s co-fractionate with LDs in density-gradient subcellular fractionation. Wild-type dry, stratified, 24 h- and 48 h-germinated seeds were subjected to subcellular fractionation on a continuous sucrose density gradient. Proteins were analyzed by immunoblotting using antibodies against NBR1, ATG8s, PEX14 (peroxisomal marker), VDAC1 (mitochondrial marker), ARF1 (Golgi marker), and OLE1 (LD marker). Proteins extracted form equal volumes of subcellular fractions were loaded on each lane. (**B**) AtNBR1 associates with LDs on two distinct structures. Hypocotyls of stratified (0), 24 h- or 48 h-germinated seeds from Arabidopsis transgenic line expressing p35S:mCherry-NBR1 (mCh-NBR1, magenta) were imaged by CLSM after LD staining with Bodipy 493/503 (green). Empty white arrowheads show puncta bearing mCh-NBR1 in close contact with LDs. Solid white arrowheads show mCh-NBR1-positive ring structures surrounding LDs (magnified views in inset). Dashed white arrowhead shows mobile aggregated LDs co-localizing with mCh-NBR1 inside the vacuole at 48 h of germination (magnified view in inset). Five images of the mobile structure acquired at different observation times are presented. (**C**) mCh-NBR1-containing structures in hypocotyls of 24 h-germinated seeds observed in (**B**) were quantified using Image J. The box plot with Tukey-style whiskers shows the number of total puncta carrying mCh-NBR1 and those carrying mCh-NBR1 surrounding LDs (n = 10 images). The red horizontal lines indicate the medians, while the bottom and top edges of the boxes represent the first (Q1) and third (Q3) quartiles, respectively. (**D**) GFP-ATG8f colocalizes with mCh-NBR1 on puncta close to LDs (empty white arrowheads) and, more rarely, on phagophore-like ring structures surrounding LDs (solid white arrowheads). Hypocotyls of 24 h-germinated seeds from Arabidopsis transgenic line co-expressing p35S:mCherry-NBR1 and pactin:GFP-ATG8f (mCh-NBR1, magenta ; GFP-ATG8f, green) were imaged by CLSM after LD staining with HCS LipidTOX 637/655 (blue). Magnified views are shown in the insets. (**E**) Magnification of the boxed region from panel **D** (upper right side). The relative intensity profile along the vector (arrow) shows superposition of GFP-ATG8f (green) and mCh-NBR1 (magenta) signals, surrounding a LD (blue). Bars = 5 µm. **<u>Alt text</u>:** Figure 2: AtNBR1 and ATG8f associate with LDs via two distinct structures during seed germination; Figure 2A: AtNBR1 and ATG8s co-fractionate with LDs in density-gradient subcellular fractionation; Figure 2B: AtNBR1 associates with LDs on two distinct structures : puncta bearing mCh-NBR1 in close contact with LDs and mCh-NBR1-positive ring structures surrounding LDs ; Figure 2C: Comparative counting of LDs surrounding by mCh-NBR1 (ring) and of puncta carrying mCh-NBR1; Figure 2D: GFP-ATG8f colocalizes with mCh-NBR1 on puncta close to LDs and, more rarely, on phagophore-like ring structures surrounding LDs ; Figure 2E: The relative intensity profile along the vector shows superposition of GFP-ATG8f and mCh-NBR1 signals, surrounding a LD.

To confirm and further investigate the association of AtNBR1 with LDs, we analyzed AtNBR1 localization by Confocale Laser Scanning Microscopy (CLSM) in transgenic Arabidopsis seeds expressing p35S:mCherry-NBR1. Hypocotyls and cotyledons from stratified (0 h-), 24 h- or 48 h-germinated seeds were imaged after neutral lipid staining with Bodipy 493/503 to visualize LDs. In both hypocotyls (Fig. 2B) and cotyledons (Fig. S1) of stratified seeds, we observed numerous large cytoplasmic puncta labeled with mCherry-NBR1 similar to the punctate structures previously reported by Svenning et al. (2011) in Arabidopsis plantlets. Notably, these structures were localized in close proximity to LDs. In 24 h-germinated seeds, we observed LDs distinctly surrounded by mCherry-NBR1, alongside the puncta previously observed in stratified seeds. These ring-like structures, which are far less abundant than the puncta (roughly ten-fold less; Fig. 2C), may correspond to autophagosomes engulfing LDs. In line with this hypothesis, we observed aggregated LDs colocalizing with mCherry-NBR1 and moving within the vacuole in hypocotyls from 48 h-germinated seeds (Fig. 2B; Video S1). These structures could result from the vacuolar degradation of AtNBR1-containing LDs and are hereafter referred to as LD remnants.

### ATG8s colocalize with AtNBR1 on LD-associated puncta and LD-surrounding autophagosome-like structures

If the AtNBR1-containing rings surrounding LDs are autophagosomes, they should also contain ATG8s, a well-established marker of autophagosomal membranes. To test this hypothesis, and given the co-fractionation of AtNBR1 and ATG8s with LDs (Fig 2A), we investigated their potential co-localization using confocal microscopy in a line co-expressing p35S:mCherry-NBR1 and pactin:GFP-ATG8f. We observed few LDs surrounded by both mCherry-NBR1 and GFP-ATG8f in 24 h-germinated seeds (Fig. 2D-E). The magnified inset in the lower right panel of Fig. 2D shows a structure resembling a phagophore engulfing a LD, potentially representing an intermediate state leading to the formation of fully closed autophagosomes around LDs. The colocalization of mCherry-NBR1 with GFP-ATG8f on a structure closely surrounding LDs was further confirmed by overlaying their fluorescence intensity profiles along a line drawn across a HCS LipidTox 637/655 stained LD (Fig. 2E). We also observed numerous puncta positive for both mCherry-NBR1 and GFP-ATG8f in close proximity to LDs (Fig. 2D). We quantified the number of these puncta per image, along with those carrying mCherry-NBR1 or GFP-ATG8f (Fig. S2). Of the mCherry-NBR1 puncta, 34 % also carried GFP-ATG8f, while 66 % of the GFP-ATG8f puncta carried mCherry-NBR1. This observation indicates that AtNBR1 and ATG8f could act together on LD dynamics, but also independently of each other. The colocalization between AtNBR1 and ATG8s on LDs observed by CLSM, together with their co-fractionation during subcellular fractionation, suggests that AtNBR1 and ATG8s could participate in the selective autophagy of a subset of LDs.

### Exploring the molecular determinants of AtNBR1 and ATG8 recruitment to LDs

AtNBR1 is characterized by the presence of a canonical ATG8-interacting motif (AIM) and two ubiquitin-associated domains (UBAs). Notably, the association of AtNBR1 with LDs coincides with the extensive ubiquitination of the major LD proteins, the oleosins, which occurs during seed germination (Deruyffelaere et al., 2015). We therefore hypothesized that AtNBR1 could act as a lipophagy receptor that binds ubiquitinated oleosins via its UBA2 domain and recruits ATG8s to LDs through its AIM domain.

To test this hypothesis, we analyzed whether the association of AtNBR1 and ATG8s with LDs was impaired in oleosin-deficient germinating seeds. Two Arabidopsis lines were analyzed: the *ole1* single mutant lacking the major oleosin OLE1, and the *ole1,2,3,4* quadruple mutant deficient in four of the five seed oleosins and retaining only the quantitatively minor oleosin OLE5. Immunoblot analysis of LD proteins revealed that AtNBR1 and ATG8s still associate with LDs in oleosin-deficient seeds (Fig. 3A), indicating that oleosins are not essential for AtNBR1 and ATG8s recruitment to LDs. Nevertheless, this result does not rule out ubiquitinated oleosins as LD-anchoring sites and cargoes for AtNBR1, alongside other LD proteins. Under this assumption, NBR1 deficiency would be expected to reduce the degradation of ubiquitinated oleosins. To explore this hypothesis, we analyzed ubiquitinated oleosin levels in two *nbr1* mutants, the *nbr1*-KO1 mutant previously generated by Tarnowski et al. (2020a) and one we isolated, named *nbr1-5*. These mutants were selected from a set of five based on the complete absence of full-length AtNBR1 detected by immunoblotting (Fig. S3). The abundance of oleosins conjugated to K48-linked diubiquitin (K48Ub₂) or K63-linked diubiquitin (K63Ub₂), the predominant ubiquitinated oleosins in germinating seeds (Deruyffelaere et al., 2015), was analyzed in these two mutants. Immunoblot analyses of LDs purified from 48 h-germinated seeds showed unchanged levels of K48Ub₂- or K63Ub₂-conjugated oleosins between the wild type and the *nbr1* mutants (Fig. 3B). This observation is in line with the one reported above, confirming that ubiquitinated oleosins are unlikely to constitute the primary LD docking sites and cargoes of AtNBR1.

**Figure 3.**
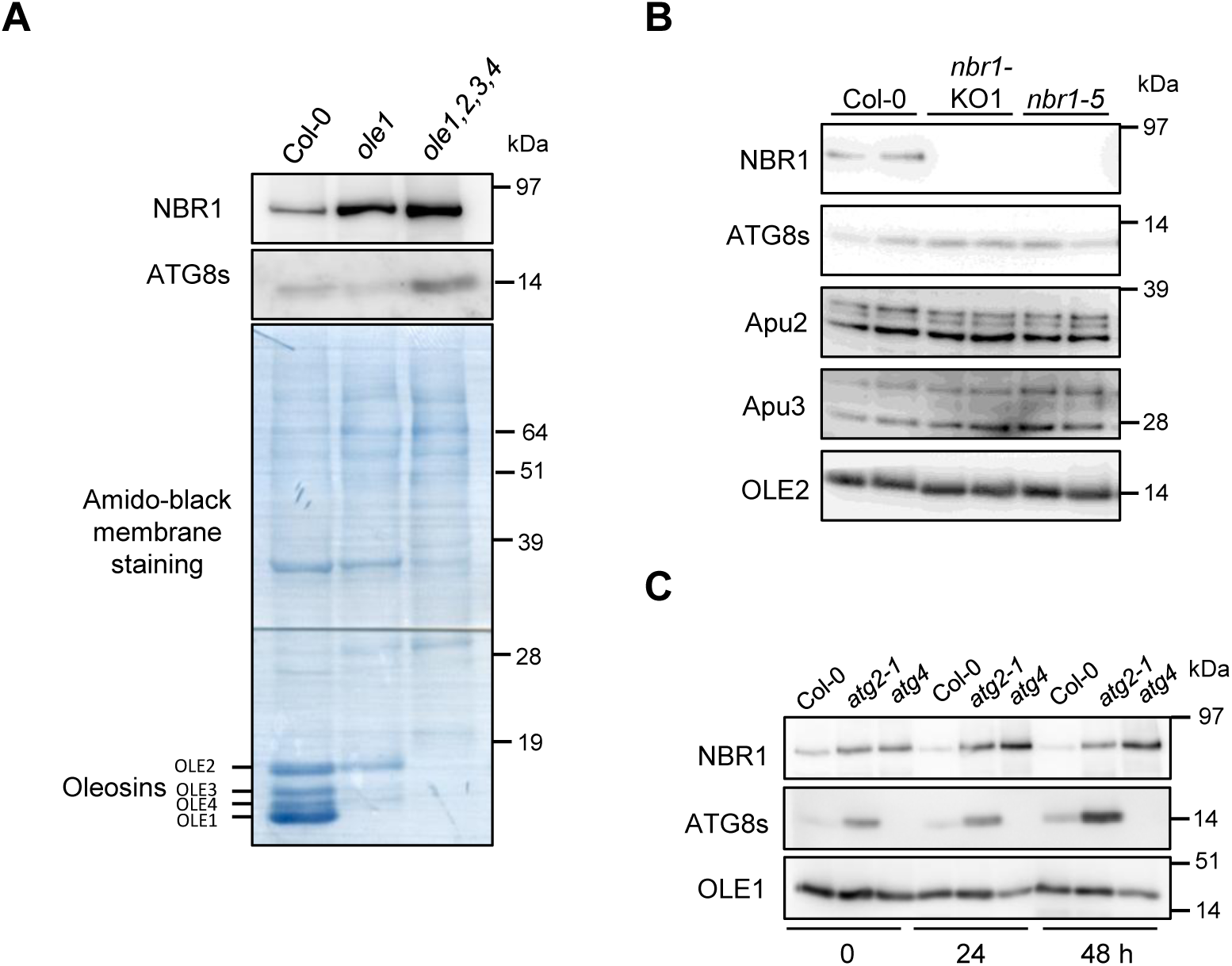
Exploring the molecular determinants of AtNBR1 and ATG8s targeting to LDs. (**A**) The association of AtNBR1 and ATG8s with LDs is not dependent on oleosins. LDs were purified from 24 h-germinated seeds from wild-type (Col-0) or *oleosin* mutants (*ole1* or *ole1,2,3,4*) and analyzed by immunoblotting. Protein profiles were visualized by amido-black staining of the membrane. (**B**) The association of ATG8s with LDs is not strictly dependent on AtNBR1. LDs were purified from 48 h-germinated seeds of wild-type (Col-0), *nbr1*-KO1 or *nbr1-5* mutants (n = 2 independent plants each) and analyzed by immunoblotting. Apu2 and Apu3 antibodies revealed K48Ub₂- and K63Ub₂-conjugated oleosins, respectively. (**C**) The association of AtNBR1 with LDs is not dependent on ATG8s. LDs were purified from stratified (0 h) or germinated seeds (24 h and 48 h) of wild-type (Col-0) or autophagy mutants (double mutant *atg4a4b-1* (*atg4*), *atg2-1)* and analyzed by immunoblotting. For all immunoblotting experiments, proteins were analyzed with the specified antibodies. Equal amounts of LD proteins were loaded onto the gel and verified with anti-OLE1 or anti-OLE2 antibodies. **<u>Alt text</u>:** Figure 3: Exploring the molecular determinants of AtNBR1 and ATG8s targeting to LDs; Figure 3A: The association of AtNBR1 and ATG8s with LDs is not dependent on oleosins; Figure 3B: The association of ATG8s with LDs is not strictly dependent on AtNBR1; Figure 3C: The association of AtNBR1 with LDs is not dependent on ATG8s.

We next assessed whether ATG8 recruitment to LDs depends on AtNBR1, and vice versa. The association of ATG8s with LDs was unchanged in the absence of AtNBR1 (Fig. 3B). Conversely, AtNBR1 association with LDs was maintained in the autophagy-deficient mutants *atg4a4b-1* (hereafter *atg4*) and *atg2-1* (Fig.3C). This association was even higher than in the wild type, in line with the well-documented overaccumulation of NBR1 and ATG8s in autophagy-deficient backgrounds (Svenning et al., 2011; Young et al., 2019; Hashimi et al., 2021). Strikingly, AtNBR1 remained associated with LDs in the *atg4* mutant, whereas ATG8s did not. This suggests that ATG8 lipidation, blocked in the *atg4* mutant, is essential for ATG8s to associate with LDs. Overall, these observations indicates that AtNBR1 and ATG8s can be recruited to LDs at least partially independently, consistent with microscopic observations shown above.

### Autophagy plays a minor role in AtNBR1-associated LD turnover

To test the hypothesis that AtNBR1 acts as a lipophagy receptor, we examined whether autophagy contributes to the degradation of AtNBR1 and LDs. To this end, we assessed the impact of pharmacological modulation or genetic disruption of autophagy on AtNBR1 and LD fates.

As a first step, we investigated whether autophagy activation using the TOR inhibitor AZD-8055 (AZD) or inhibition of vacuolar degradation with concanamycin A (ConA) enhanced the vacuolar accumulation of LDs and AtNBR1, which was occasionally observed under basal conditions as shown in Fig. 2B. The effects of these drugs were analyzed by CLSM in seedlings expressing mCherry-NBR1 (Fig 4A; Video S2). In DMSO-treated control, we observed a few mCherry-NBR1 puncta mobile within the vacuole, with LDs sporadically associating with them. AZD or ConA treatment alone did not elicit any additional effect. However, the combined application triggered the formation of aggregated mCherry-NBR1-positive LDs, which were mobile within the vacuole. To verify that these structures were indeed vacuolar, we analyzed seedlings co-expressing mCherry-NBR1 and the tonoplast marker line vac-gk (Nelson et al., 2007). The LD remnants associated with mCherry-NBR1 were confirmed to move within the tonoplast-delineated vacuole (Fig. S4; Video S3). These observations support the existence of a minor lipophagic pathway, likely mediated by AtNBR1, which is revealed more clearly upon autophagy activation and inhibition of vacuolar degradation using drugs.

**Figure 4.**
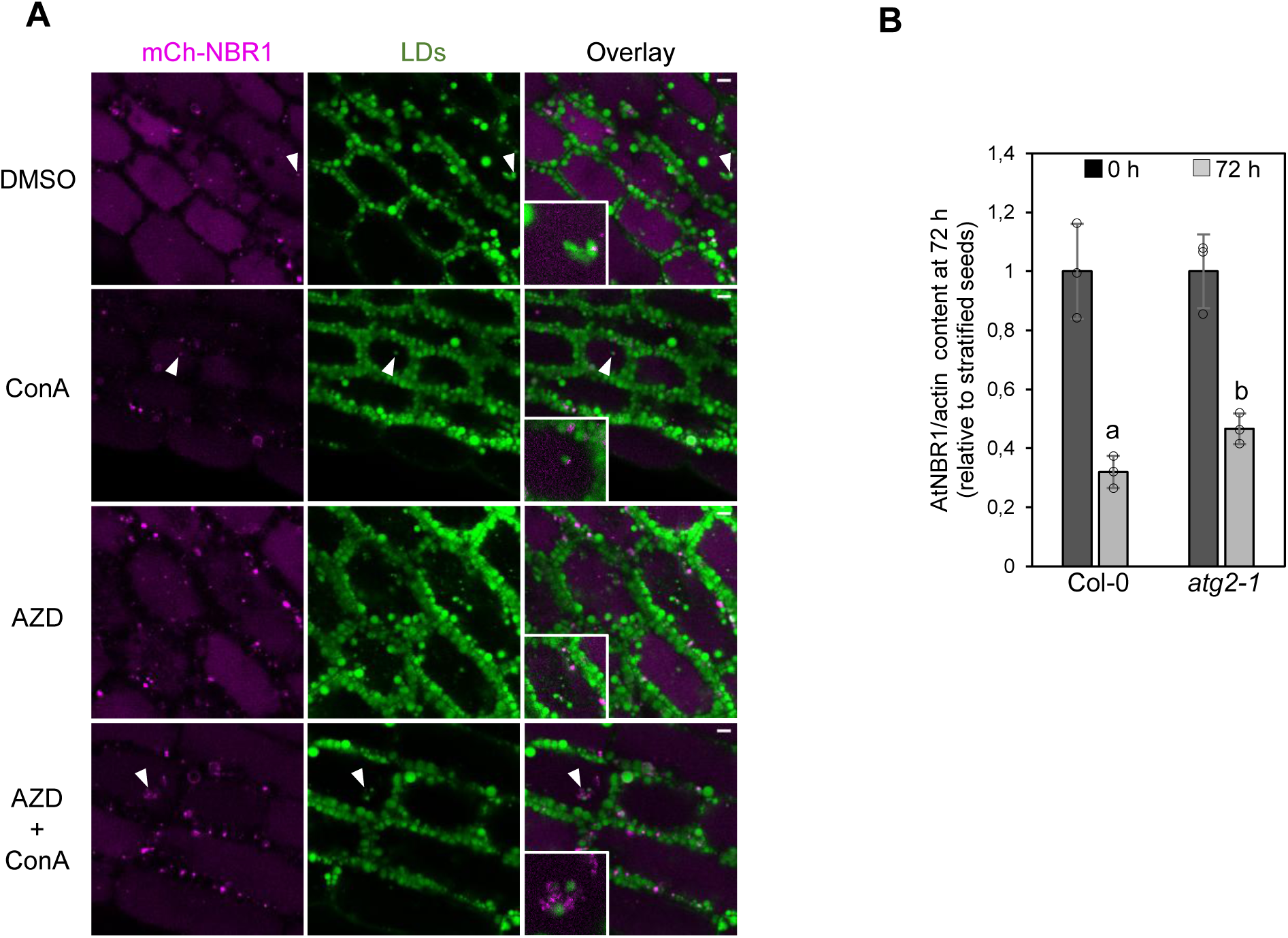
Impact of autophagy on AtNBR1 and LD turnover and fate. (**A**) Effect of autophagy modulation on the presence of mCherry-NBR1-positive LD remnants within the vacuole. Hypocotyls of 42 h-germinated seeds from Arabidopsis transgenic line expressing p35S:mCherry-NBR1 (mCh-NBR1, magenta) were treated for 5 h with DMSO, ConA, AZD or both and imaged by confocal microscopy after LD staining with Bodipy 493/503 (green). Solid white arrowheads show mobile LDs colocalizing with mCh-NBR1 inside the vacuole (magnified view in inset; Supplementary video S2). Bars = 5 µm. (**B**) A subset of AtNBR1 requires the autophagy pathway for degradation. Proteins were extracted from stratified (0 h) or 72 h-germinated seeds of Col-0 or *atg2-1* mutant and analyzed by immunoblotting with anti-NBR1 and anti-actin antibodies. AtNBR1 content normalized to actin was measured using quantitative immunoblot (mean +/- SD from n = 3 samples of 50 seeds from independent plants each). Significant difference from stratified seeds was estimated using an exact one-tailed Mann-Whitney U Test (p=0.05). Individual values are represented by open circles, and significant differences are indicated by different letters. **<u>Alt text</u>:** Figure 4: Impact of autophagy on AtNBR1 and LD turnover and fate; Figure 4A: Combined AZD + ConA treatment induces accumulation of mCherry-NBR1-positive LD remnants within the vacuole; Figure 4B: AtNBR1 degradation was delayed in the autophagy-deficient *atg2-1* mutant compared with Col-0 after 72 h of germination.

To minimize drug side effects and potential indirect responses, we subsequently analyzed AtNBR1 degradation in autophagy-deficient seeds. Total proteins from stratified (0 h) and 72 h-germinated seeds of the *atg2-1* mutant and its wild-type background were analyzed by immunoblotting to assess AtNBR1 content relative to actin (Fig. 4B). AtNBR1 abundance decreased upon germination in both genotypes, but this decrease was significantly less pronounced in the *atg2-1* mutant than in Col-0. This result indicates that AtNBR1 degradation still occurs even when autophagy is impaired, but a fraction of AtNBR1 requires the autophagy pathway for clearance, supporting its role in selective autophagy in seedlings. Together with our previous results, these observations suggest that AtNBR1 undergoes autophagic degradation and may contribute to lipophagy, although this is not the main pathway driving AtNBR1 and LD turnover in young seedlings.

### Proteasome inhibition induces AtNBR1 accumulation during germination

Since autophagy does not appear to be the primary pathway for AtNBR1 degradation, we shifted our focus to the other major protein degradation pathway in eukaryotes, the UPS. Firstly, we assessed the impact of proteasomal inhibition by MG132 on AtNBR1 levels during seed germination. MG132 treatment increased AtNBR1 level at both 24 and 48 h of germination (Fig. 5A). However, whether this observation results from enhanced *NBR1* transcription, impaired degradation, or both remained unclear. To address this, we analyzed *NBR1* transcript levels of wild-type seeds germinated for 24 h in the presence of MG132 or DMSO (Fig. 5B). MG132 significantly increased *NBR1* transcript levels, indicating that AtNBR1 accumulation under proteasome inhibition is at least partly transcriptionally regulated in germinating seeds. However, this does not exclude the possibility that MG132 also promotes AtNBR1 accumulation by inhibiting its proteasomal degradation. To test this hypothesis, we inhibited translation with cycloheximide to block MG132-induced AtNBR1 synthesis. Under these conditions, MG132 no longer induced AtNBR1 accumulation (Figure S5), suggesting that AtNBR1 degradation is proteasome-independent. However, cycloheximide severely impaired post-germinative growth. Thus, the lack of MG132-induced AtNBR1 accumulation in cycloheximide-treated seeds is difficult to interpret, as cycloheximide exerts broad systemic effects that may affect degradation pathways. We next asked whether MG132-induced AtNBR1 associate with LDs, by analyzing AtNBR1 levels in the LD fraction purified from germinated seeds treated with MG132. This treatment increased NBR1 levels to a similar extent in the total extract and the LD fraction, indicating that the upregulated AtNBR1 is associated with LDs (Fig. 5C). Overall, while these results do not provide definitive insight into the role of the proteasome in AtNBR1 degradation, the MG132-induced accumulation of AtNBR1 at LDs suggests a role for LDs in maintaining protein homeostasis during seed germination.

**Figure 5.**
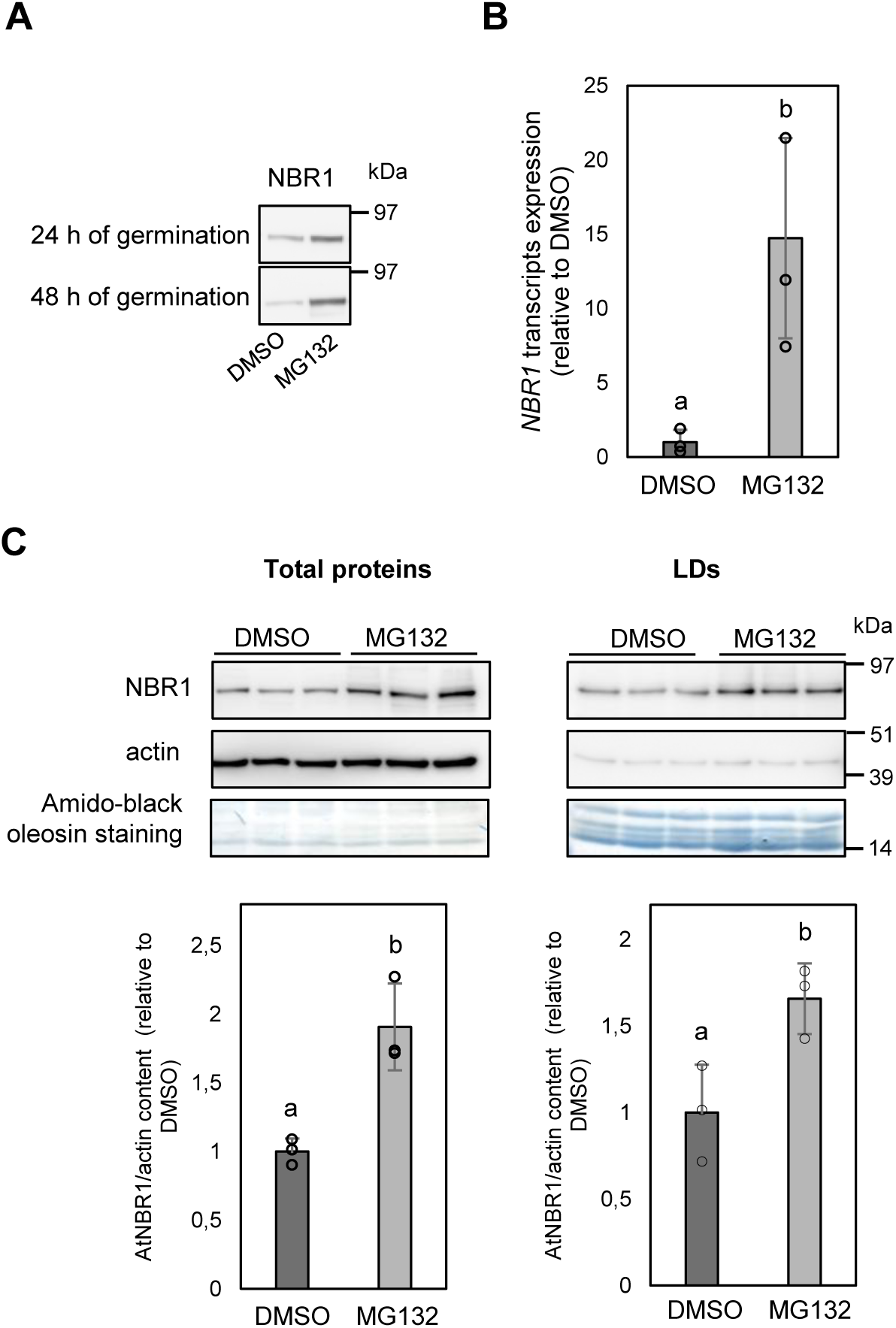
Proteasome inhibition induces AtNBR1 accumulation during germination. (A) MG132 induces AtNBR1 accumulation. Proteins were extracted from wild-type seeds germinated for 24 or 48 h in water containing 100 µM MG132 or DMSO (solvent control) and analyzed by anti-NBR1 immunoblotting. (**B**) AtNBR1 accumulation in MG132-treated seeds is transcriptionally regulated. *NBR1* expression in wild-type seeds germinated for 24 h in water containing 100 µM MG132 or DMSO (solvent control) was analyzed by RT-qPCR. *NBR1* transcript levels were normalized using *ASAR1* and *MAP2B* as reference genes (mean +/- SD from n = 3 samples of 100 seeds from independent plants). (**C**) AtNBR1 accumulated in response to MG132 associates with LDs. Wild-type seeds from three different plants were germinated for 24 h then incubated for 17 h in water containing 100 µM MG132 or DMSO. Total or LD proteins were analyzed by immunoblotting to quantify AtNBR1 levels normalized to actin (mean +/- SD from n = 3 samples of 50 seeds from independent plants). For panel (**C)**, equal loading of total proteins was verified with anti-actin antibodies, and of LD proteins by amido black staining. In panels (**B)** and (**C)**, significant difference from DMSO control was estimated using an exact one-tailed Mann-Whitney U Test (p=0.05). Individual values are represented by open circles, and significant differences are indicated by different letters. **<u>Alt text</u>:** Figure 5: Proteasome inhibition induces AtNBR1 accumulation during germination; Figure 5A: Proteasome inhibition by MG132 induces AtNBR1 accumulation in germinated seeds; Figure 5B: Proteasome inhibition by MG132 increases *NBR1* transcript levels in seeds after 24 h of germination; Figure 5C: AtNBR1 accumulated in response to MG132 associates with LDs.

### LDAD pathway contributes to the degradation of LD-associated AtNBR1

To further explore the contribution of the proteasome to the degradation of LD-associated AtNBR1, we investigated the potential involvement of the LDAD pathway, which mediates the extraction of proteins from the LD surface for subsequent proteasomal processing (Deruyffelaere et al., 2018; Kretzschmar et al., 2018). The core components of this pathway are PUX10, which recognizes ubiquitinated LD proteins, and the segregase CDC48A which mediates their removal from LDs for proteasomal degradation. We assessed the contribution of LDAD to AtNBR1 turnover by analyzing *pux10* mutants. We explored the degradation of AtNBR1 in dry, stratified (0 h) and germinated seeds (24 h to 72 h) of Col-0 and *pux10* mutants (Fig. 6A). AtNBR1 degradation was delayed during seed germination in the *pux10-1* mutant, while its accumulation during stratification remained unchanged. A similar result was also obtained with the *pux10-2* mutant (Fig. S6). Notably, AtNBR1 levels increased in *pux10-1* without changes in *NBR1* transcript levels (Fig. 6B), indicating that AtNBR1 accumulation results from delayed degradation rather than enhanced transcription. We also demonstrated that AtNBR1 accumulating in *pux10-1* mutant associates with LDs, as shown by immunoblot analysis of LD proteins (Fig. 6C-D). Overall, this phenotype mirrors that reported for oleosins (OLE2) and their ubiquitinated forms (OLE-K48Ub_2_) (Deruyffelaere et al., 2018). These results highlight a role for PUX10 in AtNBR1 degradation.

**Figure 6.**
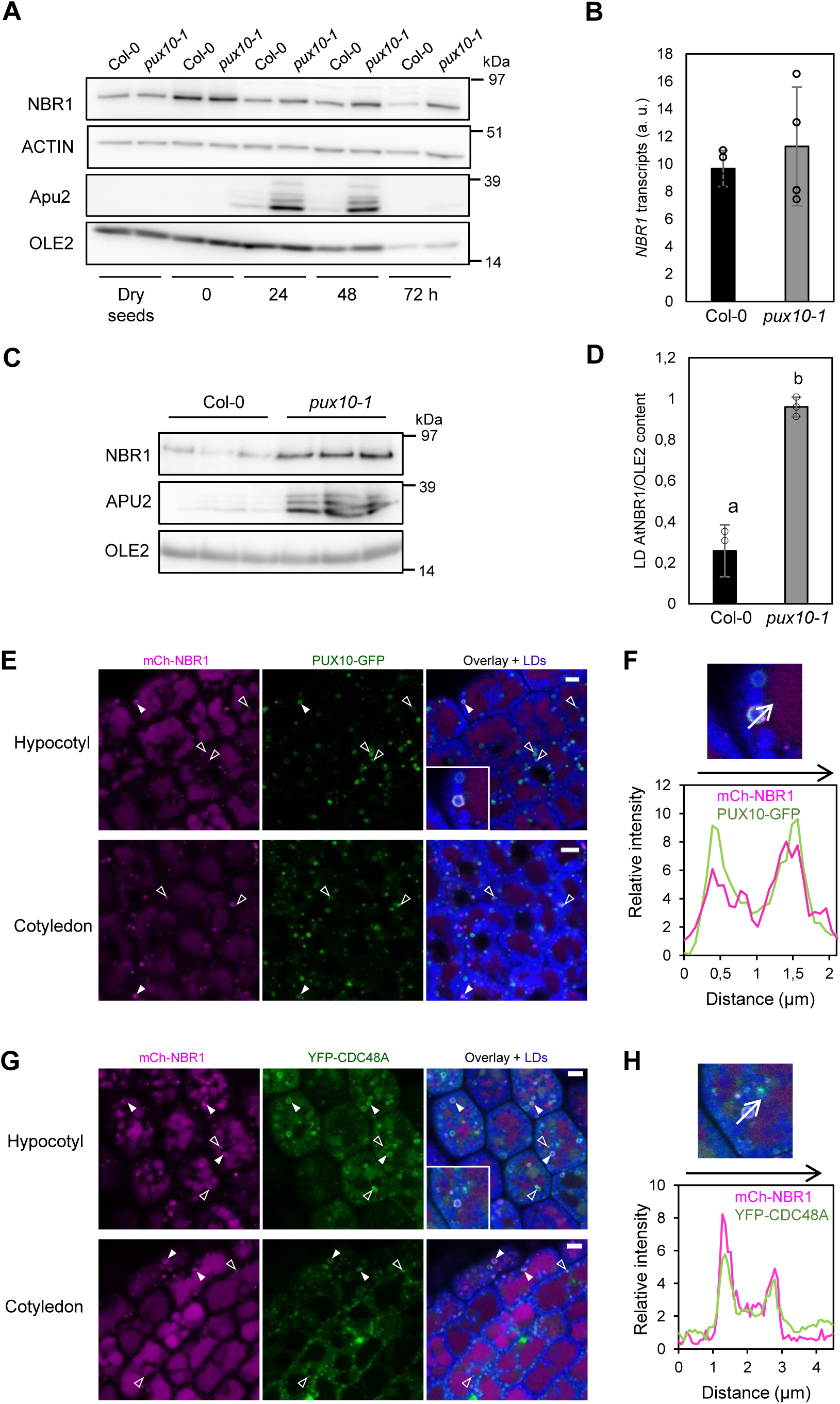
LDAD pathway contributes to AtNBR1 degradation. A) Degradation of AtNBR1 is delayed in *pux10-1* mutant. Proteins extracted from an equivalent amount of dry, stratified (0 h) or germinated seeds (24 to 72 h) of wild-type (Col-0) or *pux10-1* were analyzed by immunoblotting with the specified antibodies. **(B)** AtNBR1 is not transcriptionally regulated in *pux10* mutant. *NBR1* transcript levels were analyzed by RT–qPCR in Col-0 or *pux10-1* seeds germinated for 48 h and normalized using *ASAR1* and *MAP2B* as reference genes (mean +/- SD from n = 4 samples of 100 seeds from independent plants). (**C**) AtNBR1 accumulated in *pux10-1* mutant is associated with LDs. LDs were purified from Col-0 or *pux10-1* seeds germinated for 52 h and analyzed by immunoblotting with the specified antibodies, using OLE2 as a loading control. (**D**) AtNBR1 relative to OLE2 content from (**C**) was measured using quantitative immunoblot (mean +/- SD from n = 3 samples of 50 seeds from independent plants). (**E**) AtNBR1 and PUX10 partially colocalize on LDs. Cotyledons and hypocotyls from 24 h-germinated Arabidopsis lines co-expressing p35S:mCherry-NBR1 (mCh-NBR1, magenta) and pPUX10:PUX10-GFP (PUX10-GFP, green) were stained with HCS LipidTOX 637/655 (LDs, blue) and imaged by CLSM. Solid white arrowheads show LDs totally surrounded by both mCh-NBR1 and PUX10-GFP. Empty white arrowheads show vesicles tagged with both mCh-NBR1 and PUX10-GFP in close contact with LDs. Magnified view are shown in inset. (**F**) Magnification of boxed region from (**E**). The relative intensity profile along the vector (arrow) shows superposition of PUX10-GFP (green) and mCh-NBR1 (magenta) signals. (**G**) AtNBR1 and CDC48A partially colocalize on LDs. As in (**E**), using an Arabidopsis line co-expressing p35S:mCherry-NBR1 (mCh-NBR1, magenta) and pCDC48:YFP-CDC48A (YFP-CDC48A, green). Solid white arrowheads show LDs totally surrounded by both mCh-NBR1 and YFP-CDC48A. Empty white arrowheads show vesicles tagged with both mCh-NBR1 and YFP-CDC48A in close contact with LDs. Magnified view are shown in inset. (**H**) Magnification of boxed region from (**G**). The relative intensity profile along the vector (arrow) shows superposition of YFP-CDC48A (green) and mCh-NBR1 (magenta) signals. Bars = 5 µm. In panels **B,** no statistically significant differences were detected using the Mann–Whitney U test (P > 0.05). In panel **C**, significant difference from Col-0 was estimated using an exact one-tailed Mann-Whitney U Test (p=0.05). Individual values are represented by open circles, and significant differences are indicated by different letters. **Alt text:** Figure 6: LDAD pathway contributes to AtNBR1 degradation; Figure 6A: Degradation of AtNBR1 is delayed in *pux10-1* mutant; Figure 6B: *NBR1* transcript levels are similar in Col-0 and the *pux10-1* mutant, indicating that AtNBR1 accumulation in the *pux10-1* mutant is not due to transcriptional upregulation; Figure 6C: AtNBR1 accumulated in *pux10-1* mutant is associated with LDs; Figure 6D: The NBR1 relative to OLE2 content from (C) was quantified and it showed that the increased AtNBR1 accumulation in the *pux10-1* mutant is associated with LDs; Figure 6E: AtNBR1 and PUX10 partially colocalize on LDs; Figure 6F: The relative intensity profile along the vector shows superposition of PUX10-GFP and mCh-NBR1 signals; Figure 6G: AtNBR1 and CDC48A partially colocalize on LDs; Figure 6H: The relative intensity profile along the vector shows superposition of YFP-CDC48A and mCh-NBR1 signals.

To further investigate the relationship between the LDAD pathway and AtNBR1, we examined AtNBR1 localization relative to the LDAD components PUX10 and CDC48A. To this end, transgenic seedlings co-expressing mCherry-NBR1 with either PUX10-GFP or YFP-CDC48A were imaged by CLSM. We observed partial co-localization of mCherry-NBR1 with PUX10-GFP (Fig. 6E) or YFP-CDC48A (Fig. 6G) at punctate structures adjacent to LDs. Some LDs were also distinctly surrounded by both mCherry-NBR1 and PUX10-GFP (Fig. 6E) or YFP-CDC48A (Fig. 6G). Fluorescence intensity profiles along a line drawn across one such LD confirmed overlap of the fluorescence signals for each protein pair (Fig. 6F-H).

Altogether, the partial co-localization of AtNBR1 with components of LDAD machinery, along with the delayed AtNBR1 degradation in the PUX10-deficient background, strongly supports a role for the LDAD pathway in the degradation of LD-associated AtNBR1 during seed germination.

### NBR1 is not involved in a major LD degradation pathway

To further investigate the contribution of AtNBR1 to LD dynamics, we compared fatty acid content of dry, stratified (0 h) and germinated seeds from Col-0 and two independent *nbr1* mutant lines. At all stages, no significant differences were detected between the genotypes, either in total fatty acid content (Fig. S7 A) or in fatty acid composition (data not shown). These findings indicate that AtNBR1 plays, at most, a minor role in fatty acid mobilization during germination. To further support this conclusion, we assessed the LD-integral protein degradation in both wild-type and *nbr1* mutant lines. Total proteins extracted from 48 h-germinated seeds of the Col-0, *nbr1*-KO1 and *nbr1-5* mutant lines were analyzed by immunoblotting using antibodies directed against LD-integral proteins (Fig. S7 B). No differences were detected among the lines. We additionally quantified OLE1 degradation by immunoblotting in 48 h-germinated seeds of *nbr1* mutants and Col-0 (Fig. S7 C). Again, no detectable differences were observed, further supporting the conclusion that AtNBR1 is not involved in a major LD degradation pathway.

### Predominance of proteostasis-related proteins in the AtNBR1-dependent LD-associated proteome

To learn more about the role of LD-associated AtNBR1, we sought to identify its binding partners using two complementary approaches, focusing on the stratified seed stage, since it corresponds to maximal AtNBR1 accumulation at LDs, as reported above (Fig. 1 and 2).

In one approach, we performed mCherry pull-down experiments on purified LDs from p35S:mCherry-NBR1 and Col-0 stratified seeds. Eluates from RFP-trap beads were analyzed by LC-MS/MS, resulting in the identification of 680 proteins with at least two unique peptides per protein (Supplementary Data Set Table S1 A). After normalization, spectral counts from mCherry-NBR1 LD pull-downs were compared with those of Col-0 LDs. Twenty proteins were either at least twofold significantly enriched (p<0.05) or exclusively detected in LDs containing mCherry-NBR1, identifying them as potential binding partners of AtNBR1. This dataset was significantly enriched (p<0.05) in proteins associated with the GO molecular function terms “mRNA binding” and “structural constituents of ribosome” as assessed using g:GOST. Importantly, it also contains several proteins involved in protein folding that belong to the small HSPs family, such as HSP17.6A. Acting as ATP-independent first-line chaperones, these proteins, like NBR1, contribute to the PQC. Interestingly, the most strongly enriched protein, phytochrome A (PHYA), a major photoreceptor that mediates light-dependent developmental responses, including seed LD mobilization (Kozuka et al., 2020), was recently shown to interact with ATG8s and negatively regulate autophagosome formation under far-red light (Jiang et al., 2026), raising the question of whether LDs and AtNBR1 contribute to this regulation. Notably, two other proteins identified in this dataset, LIPA and TSPO, have previously been implicated in LD–plasma membrane tethering and in the regulation of LD abundance in seeds, respectively (Jurkiewicz et al., 2018; Krawczyk et al., 2022).

In the second approach, we analyzed the proteome of LDs purified from stratified seeds of Col-0 and the *nbr1*-KO1 mutant by LC-MS/MS, and, after normalization, we compared the enrichment of proteins in Col-0 LDs relative to *nbr1*-KO1 LDs. We detected a total of 856 proteins with at least two unique peptides per protein (Supplementary Data Set Table S1 B). Among them, twenty-eight proteins were significantly enriched by at least two-fold (p<0.05) or only detected in Col-0 compared with *nbr1*-KO1 line. We focused on these proteins as candidates that directly or indirectly interact with AtNBR1 on LDs. This dataset was significantly enriched (p<0.05) in proteins associated with the GO molecular function terms “unfolded protein binding” and “oxidoreductase activity”, as assessed using g:GOST. The large set of metabolic enzymes, including oxidoreductases, could reflect the metabolic reactivation that occurs upon imbibition. The biological significance of the AtNBR1-dependent association of these enzymes with LDs remains unclear but may suggest that LDs act as a reservoir for metabolic enzymes during metabolic reactivation. Like the pull-down dataset, this dataset contains proteins involved in PQC, including proteins that regulate protein synthesis and folding, as well as components related to the UPS degradation pathway. Notably, we identified RAD23D, that was absent from *nbr1-*KO1 LDs. RAD23D is a member of the RADIATION SENSITIVE23 family, whose members function as ubiquitin receptors that shuttle ubiquitinated substrates to the 26S proteasome for degradation in Arabidopsis (Farmer et al., 2010). Interestingly, a comparable function has been attributed to NBR1 in mammals, where p62/SQSTM1 has been proposed to act as a shuttling factor that delivers ubiquitinated proteins to the proteasome via its PB1 and UBA2 domains (Seibenhener et al., 2004a; Seibenhener et al., 2004b; Geetha et al., 2008). In addition, in yeast, Cdc48 has been shown to participate in the Ufd2/Rad23-dependent proteolytic process known as the “Ubiquitin Fusion Degradation” pathway. In this system, Cdc48 is thought to release Rad23 from its interactor Ufd2, thereby enabling its association with Rpn1, a proteasome subunit (Richly et al., 2005; Baek et al., 2011).

By integrating the results of the two approaches (Table S1), we identified a single protein that was significantly enriched both in LDs from Col-0 relative to *nbr1*-KO1 and in the pull-down of LDs from mCherry-NBR1 relative to Col-0. This protein, the tripeptidyl peptidase II (TPP2), forms an oligomeric exopeptidase complex involved in amino acid recycling (Book et al., 2005) and is a component of the proteasome-associated proteolytic pathway in *A. thaliana* (Polge et al., 2009). The potential interaction—whether direct or indirect—of AtNBR1 with TPP2 warrants further study but, together, these findings suggest that AtNBR1 may contribute to PQC processes on LDs.

### NBR1 contributes to seed longevity

AtNBR1 interactome analyses revealed a predominance of proteins involved in protein synthesis, folding and degradation, suggesting that AtNBR1 may play a role in LD-associated PQC. Such a role could be particularly important during seed imbibition and germination, when proteostasis is essential for seedling establishment. During seed storage, oxidative protein damage progressively accumulates, eventually overwhelming the repair capacity upon imbibition, resulting in a progressive loss of seed vigor and germination capacity (Sano et al., 2016). This process, known as seed ageing, can be experimentally reproduced by controlled deterioration treatment, also referred to as accelerated ageing (AA), which recapitulates the molecular and biochemical changes associated with natural seed ageing (Rajjou et al., 2008). This experimental system was used to investigate the role of AtNBR1 in seed longevity.

Wild-type Col-0, *nbr1-5* and *nbr1*-KO1 mutant seeds were subjected to an AA in a closed tank maintained at 40 °C and 85 % relative humidity for three to ten days. Following the treatment, seeds were germinated for eight days alongside untreated control seeds (Fig. 7). Notably, starting at five days of AA, a decline in germination capacity was observed in mutant lines compared to the wild-type. This reduction was particularly pronounced in the *nbr1*-KO1 mutant, where germination capacity was reduced by 50 % after seven days of treatment. Following ten days of AA, all lines completely lost their ability to germinate. These findings indicate that the absence of AtNBR1 significantly compromises seed longevity.

**Figure 7.**
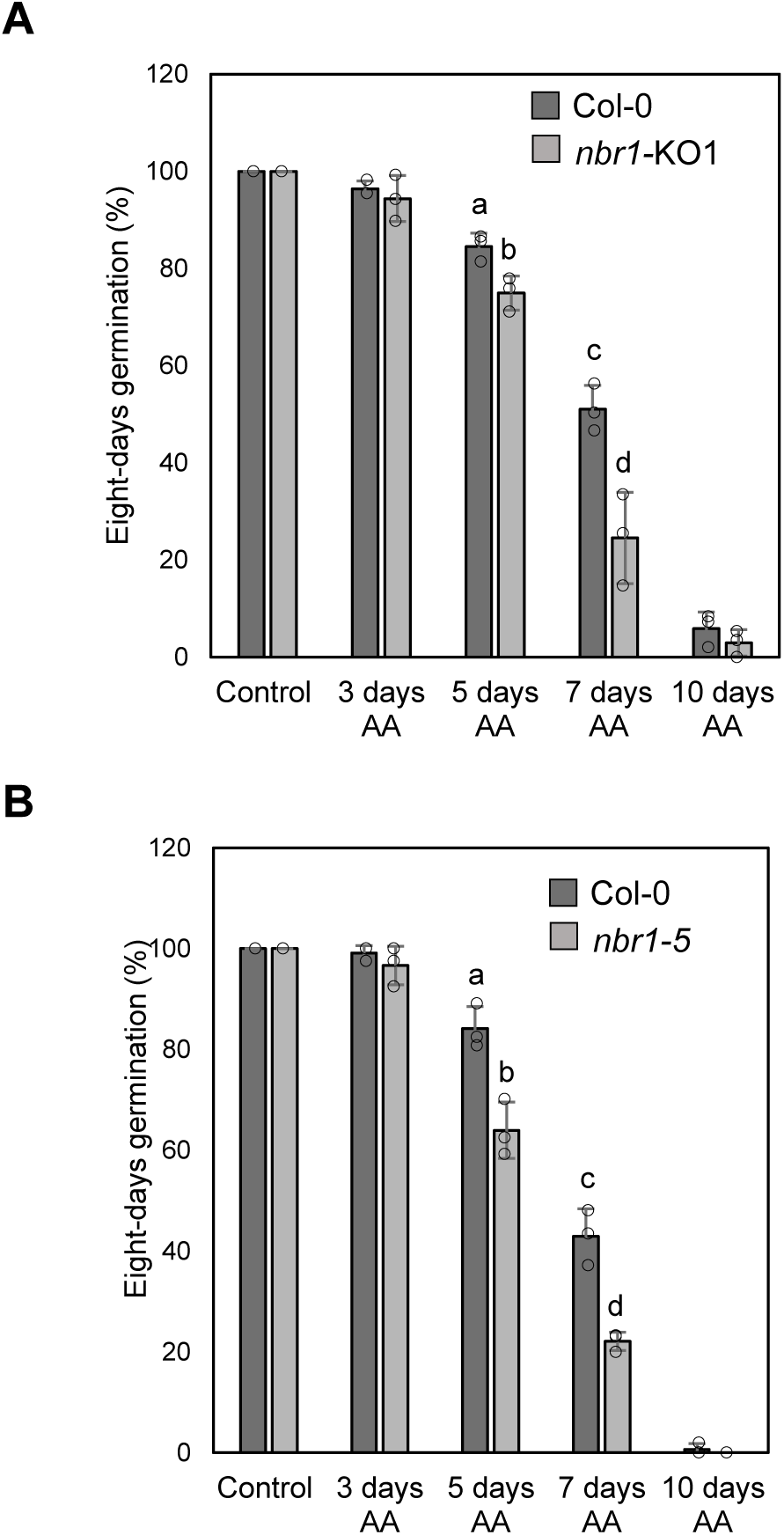
AtNBR1 contributes to seed longevity. Defect in AtNBR1 affects the seed germination capacity after accelerating aging. (**A**) Wild-type (Col-0) or *nbr1-KO1* mutant seeds underwent an accelerating aging (AA) for three to ten days, and were germinated for eight days along with control (no AA). Shown are averages of three technical repetitions of three biological replicates (mean +/- SD from n = 3 samples of 100 seeds / batch). Significant difference from Col-0 was estimated using an exact one-tailed Mann-Whitney U Test (p=0.05). Individual values are represented by open circles, and significant differences are indicated by different letters. **(B)** Same experiment as in (**A**) using another *nbr1* mutant (*nbr1-5*) compared to Col-0. **<u>Alt text</u>:** Figure 7: AtNBR1 contributes to seed longevity; Figure 7A: Defect in AtNBR1 in *nbr1*-KO1 mutant affects the seed germination capacity after accelerating aging; Figure 7B: Defect in AtNBR1 in *nbr1-5* mutant affects the seed germination capacity after accelerating aging.

## Discussion

Lipophagy remains poorly characterized in plants and largely unexplored in seeds despite their high LD content (Xu and Fan, 2022). While microlipophagy has been implicated in LD breakdown during seed germination (Poxleitner et al., 2006; Miklaszewska et al., 2023), macrolipophagy has not yet been demonstrated, and the involvement of SARs in seed LD degradation remains to be established. Our initial observation that NBR1, the best-characterized plant SAR, predominantly localizes to LDs in germinating seeds prompted us to investigate its dynamics and function during germination. While our results indicate that NBR1-associated lipophagy contributes only marginally to bulk LD mobilization, they uncover a previously unrecognized association of plant NBR1 with PQC actors at the LD surface which may underlie its effect on seed longevity. Furthermore, they identify LDAD- and UPS-related factors as potential components of an NBR1-associated proteostasis network.

### AtNBR1 and ATG8s predominantly associate with LDs in germinating seeds

In this study, we demonstrate that AtNBR1 predominantly associates with LDs during seed germination, with this association being transient. Cold imbibition triggered *NBR1* expression and AtNBR1 accumulation at LDs, followed by its progressive degradation during germination. Notably, ATG8s exhibited LD-association dynamics similar to those of AtNBR1 and co-localized with AtNBR1 in close proximity to LDs. The nearly exclusive association of AtNBR1 and ATG8s with LDs appears to be a distinctive feature of germinating seeds, as it is lost during post-germinative growth and has not been reported in other plant tissues. This suggests a functional interplay between autophagy and LDs that may be particularly important during seed germination.

NBR1 is only the third SAR identified as associating with LDs in plants. Indeed, the plant-specific ATG8-interacting proteins ATI1 and ATI2 were previously shown to relocalize from the ER to spherical structures in response to carbon starvation (Honig et al., 2012). These structures were recently identified as LDs (Li et al., 2025). In mammals, several SARs localize to LDs, as recently reviewed by Zhao et al. (2026) including SQSTM1/p62 and NBR1, the closest homologs of plant NBR1 (Robichaud et al., 2021; Shroff and Nazarko, 2023). NBR1 homologs also associate with LDs in other metazoans, including *Caenorhabditis elegans* and *Drosophila melanogaster* (Yan et al., 2019; Kumar et al., 2023). However, the molecular mechanisms underlying SAR recruitment to LDs remain poorly understood, despite being central to deciphering how the selective autophagy machinery recognizes LDs. Among studies reporting p62/SQSTM1 or NBR1 localization to LDs, only one identified an LD-associated protein mediating this association. In hepatocytes, SQSTM1/p62 recruitment to LDs during ethanol-induced lipophagy depends on the LD protein perilipin 1 (Wang et al., 2017), which can be ubiquitinated (Ju et al., 2019), suggesting a potential role for ubiquitin recognition in this process. Here, we investigated how AtNBR1 and ATG8s associate with LDs in Arabidopsis seeds. We speculate that AtNBR1 could bind ubiquitinated LD proteins through its UBA2 domain thereby promoting ATG8 association with LDs via its AIM (Svenning et al., 2011). Because oleosins, the major LD proteins, are extensively ubiquitinated during germination (Deruyffelaere et al., 2015), ubiquitinated oleosins could potentially serve as AtNBR1 cargoes to mediate its association with LDs. However, this hypothesis is not supported, as AtNBR1 recruitment to LDs was unaffected in oleosin-deficient lines and ubiquitinated oleosin level was unchanged in *nbr1* mutants compared with the wild type. Alternatively, ATG8s could undergo lipidation directly on LDs, as recently demonstrated in starved adipocytes (Omrane et al., 2023), thereby anchoring ATG8s to the LD surface and potentially promoting NBR1 recruitment. However, this model cannot account for AtNBR1 association with LDs, which persists in the *atg4* mutant despite impaired ATG8 lipidation. Conversely, the presence of ATG8s on LDs in the absence of AtNBR1 does not preclude AtNBR1 from acting upstream of ATG8, as multiple recruitment pathways involving OLE1, CLO1, ATI1/2 and AtNBR1 may coexist. Indeed, ATG8s interact with the LD proteins OLE1 and CLO1 through their LIR/AIM docking site (Marshall et al., 2019; Miklaszewska et al., 2023), while ATI1/2 are ATG8-interacting proteins, as their name indicates (Honig et al., 2012). Together, these results support a model in which AtNBR1 is recruited to LDs upstream of ATG8s, independently of the major seed LD proteins, oleosins. Future studies will be needed to identify the mechanisms driving the recruitment of AtNBR1 and ATG8s to LDs and to determine why only a subset of LDs is selectively targeted.

### AtNBR1 and ATG8s define two seed LD-associated structures

Analysis of AtNBR1 and ATG8 localization by microscopy in 24 h-germinated seeds revealed two distinct colocalization patterns: a predominant punctate pattern closely associated with LDs and a second pattern consisting of ring-like structures surrounding LDs.

The punctate foci are reminiscent of the AtNBR1-containing cytoplasmic puncta previously observed in Arabidopsis seedlings at later developmental stages (Svenning et al., 2011; Ji et al., 2020) and during recovery from heat stress (Thirumalaikumar et al., 2020). The formation of these structures relies on PB1 domain-mediated self-polymerization of AtNBR1 (Svenning et al., 2011), which drives the formation of membraneless condensates through liquid–liquid phase separation (Yan et al., 2024). These condensates resemble those formed by mammalian SQSTM1/p62, which sequester ubiquitinated protein aggregates (Danieli and Martens, 2018; Zaffagnini et al., 2018). In Arabidopsis protoplasts, AtNBR1 condensate formation precedes the recruitment of both ATG8 and the phagophore assembly site (Yan et al., 2024). Thus, the AtNBR1 puncta observed in close proximity to LDs in seeds may correspond to NBR1 condensates formed through self-polymerization, and their formation may precede ATG8 recruitment, as discussed above. Whether AtNBR1 condensates contain protein aggregates sequestered in the vicinity of LDs remains unresolved, as the differential centrifugation approach commonly used to isolate protein aggregates is compromised by their potential association with low-density LDs.

The ring-like structure surrounding LDs observed in the second AtNBR1- and ATG8-positive pattern is consistent with an autophagosome engulfing LDs. Although not definitive on its own, this observation strongly supports the existence of an AtNBR1-associated lipophagic pathway in the seed, in which AtNBR1 may function as a SAR mediating LD recognition and sequestration.

### Evidence for a minor macrolipophagic pathway involving AtNBR1 in germinated seeds

In our study, CLSM revealed not only structures resembling autophagosomes surrounding LDs and carrying both AtNBR1 and ATG8f, but also AtNBR1-associated LD remnants moving within the vacuole, consistent with the delivery of AtNBR1-positive LDs to the vacuolar lumen. Previous studies in plants have implicated autophagy-related mechanisms in non-seed LD turnover, including ATG8-associated processes and vacuolar degradation pathways (Kurusu et al., 2014; Fan et al., 2019; Xu and Fan, 2022; Qin et al., 2023). In germinated seeds, ATG8b was also reported to co-localize with LDs, although no autophagosomal membrane surrounding LDs was observed, leading to the proposal of a microlipophagy mechanism involving interactions between ATG8 and CLO1 (Miklaszewska et al., 2023). Our microscopy observations suggest that AtNBR1 may be involved in a macrolipophagy-like mechanism during seed germination, which could coexist with the microlipophagy process. Consistent with this hypothesis, the accumulation of AtNBR1-positive LD structures within the vacuole upon combined TOR inhibition and blockade of vacuolar degradation suggests that AtNBR1-associated LD material is delivered to, and rapidly degraded within, the vacuole. The absence of a major effect of AZD alone further supports the idea that these structures are highly transient and undergo rapid vacuolar turnover. In addition, the modest but significant stabilization of AtNBR1 in the *atg2* mutant indicates that a fraction of AtNBR1 undergoes constitutive autophagic degradation, pointing to a contribution of the canonical macroautophagy machinery to this process. Importantly, however, our data indicate that this pathway makes only a minor contribution to overall LD turnover during seed germination. Loss of AtNBR1 did not significantly affect lipid remobilization, consistent with lipolysis being the predominant mechanism driving LD mobilization during germination (Kelly et al., 2011). Likewise, the absence of a detectable effect of AtNBR1 loss on the degradation of integral LD proteins after 48 h of germination further supports a limited contribution of NBR1-dependent autophagy to LD turnover. Altogether, these results provide evidence for a macrolipophagic process in Arabidopsis, a mechanism partially characterized in mammals (reviewed by Zhao et al (2026)) but not yet demonstrated in plants.

Several lines of evidence support a role for AtNBR1 as a SAR in this process: AtNBR1 colocalizes with ATG8 in ring-like structures surrounding LDs, accumulates in LD remnants within the vacuole upon AZD/ConA treatment, and exhibits reduced turnover in the *atg2* mutant. However, establishing AtNBR1 as a bona fide SAR for LDs will require identifying the LD-associated cargo recognized by AtNBR1, determining whether this recognition depends on ubiquitination, and demonstrating that AtNBR1 is required for the selective delivery of LDs to the vacuole. Addressing these questions remains challenging because the contribution of this pathway to LD turnover appears to be minor and is likely masked by the predominant lipolytic pathway and the alternative CLO1-driven lipophagic pathway.

### AtNBR1 contributes to seed longevity, putatively through a proteostasis network involving LDAD and UPS pathways

During orthodox seed storage and subsequent imbibition, quiescent tissues progressively accumulate reactive oxygen species, leading to protein oxidation (Rajjou et al., 2008; Zinsmeister et al., 2026). These damaged proteins may form toxic aggregates that must be cleared to ensure successful germination, potentially through autophagy, which has been implicated in seed longevity in Arabidopsis (Minina et al., 2018; Shinozaki et al., 2024). In this context, the contribution of AtNBR1 to seed longevity observed in the present study may be linked to its established role in aggrephagy (Svenning et al., 2011). In addition to its potential role in the autophagic clearance of protein aggregates, AtNBR1 has been reported to modulate ABA signaling during seed imbibition (Tarnowski et al., 2020b), suggesting that changes in ABA sensitivity may also influence germination performance following seed aging. Nevertheless, our biochemical and proteomic data point to an alternative mechanism underlying the contribution of AtNBR1 to seed longevity, implicating UPS-related factors and highlighting a potential role for LDs in this process.

Indeed, the limited contribution of autophagy to AtNBR1 degradation suggests that alternative turnover pathways, including the UPS, may be involved. In this context, we identified the exopeptidase TPPII as a putative AtNBR1 interactor among LD-associated proteins using two independent approaches. This protease is functionally linked to the UPS (Polge et al., 2009) and has been shown to act downstream of the proteasome in mammals (Reits et al., 2004). These approaches also revealed several additional components of the PQC machinery, involved in protein synthesis, protein refolding and protein degradation, suggesting that AtNBR1 may be associated with broader proteostasis processes. Further supporting a link between AtNBR1 and the UPS, we found that PUX10, a central component of the LDAD pathway, is involved in AtNBR1 degradation. Indeed, PUX10 and CDC48A, core components of the LDAD pathway, act upstream of the UPS by selecting and extracting LD-anchored proteins targeted for degradation. Together, these observations support the possibility that AtNBR1 participates in proteostasis during seed germination through mechanisms extending beyond autophagy and involving the UPS. A similar relationship has recently been reported in rice, where OsNBR1 interacts with proteasome components to regulate the degradation of an auxin signaling regulator during root development (Sun et al., 2025). Such a role in maintaining proteome homeostasis may explain our finding that AtNBR1 positively contributes to seed longevity in Arabidopsis. Consistent with this observation, a similar contribution to seed longevity has been reported for the NBR1 interactor ROF1, which modulates chaperone accumulation during heat stress in plants (Bissoli et al., 2012; Thirumalaikumar et al., 2020).

Based on the findings presented in this study, we propose an integrative model in which AtNBR1 puncta associated with seed LDs may represent condensate-like structures enriched in proteins destined for degradation. This hypothesis is consistent with previous reports of similar structures involving AtNBR1 and its homologue in plant protoplasts and mammalian cells (Zaffagnini et al., 2018; Yan et al., 2024). Given that animal LDs can act as transient reservoirs for non-resident LD proteins prior to their relocalization or degradation, including histone sequestration in Drosophila (Stephenson et al., 2021) and ApoB degradation in human hepatocytes (Fujimoto and Ohsaki, 2006), we propose that LD-associated AtNBR1 puncta may similarly serve as platforms for the transient accumulation of unwanted or damaged proteins prior to their degradation. Within these structures at the surface of seed LDs, AtNBR1 may participate in a proteostasis network involving UPS and LDAD components to promote the clearance of damaged proteins, thereby contributing to seed longevity. This model may serve as a framework for future studies addressing several outstanding questions, including whether the AtNBR1 PB1 domain and LDAD contribute to AtNBR1 puncta formation and disassembly, respectively, and whether these processes are linked to the molecular mechanisms underlying seed longevity.

## MATERIALS AND METHODS

### Plant Material and growth conditions

*Arabidopsis thaliana* ecotypes Wassilewskija (Ws) and Columbia (Col-0) were obtained from the *Arabidopsis thaliana* resource center for genomics at the Institut Jean-Pierre Bourgin (http://publiclines.versailles.inrae.fr/). The T-DNA mutant lines *pux10-1* (SAIL_1187), *pux10-2* (FLAG_211E02), *atg2-1* (SALK_076727), *atg4a4b-1, nbr1-2* (GK_246H08), *nbr1-3* (SALK_144852) and *nbr1-4* (WiscDsLoxHs007_07A) were described in Deruyffelaere et al. (2018), Yoshimoto et al. (2009), Yoshimoto et al. (2004), Zhou et al. (2013), Jung et al. (2020) and Young et al. (2019), respectively. The T-DNA mutant line *nbr1-5* (GK_281A10) was obtained from the Arabidopsis Biological Resource Center (https://abrc.osu.edu) and selected to homozygosity. The CRISPR/Cas9 generated line *nbr1*-KO1 was described in Tarnowski et al. (2020). All lines were in the Col-0 genetic background, except *atg4a4b-1* and *pux10-2* which were in Ws background. The transgenic lines p35S:mCherry-NBR1, pactin:GFP-ATG8f, p35S:vac-gk, pPUX10:PUX10-GFP and pCDC48A:YFP-CDC48A were described in Svenning et al. (2011), Di Berardino et al. (2018), Nelson et al. (2007) and Deruyffelaere et al. (2018), respectively. Double transgenic lines were generated by crossing p35S:mCherry-NBR1 with pactin:GFP-ATG8f, pCDC48A:YFP-CDC48A, pPUX10:PUX10-GFP and p35S:vac-gk. Arabidopsis lines used in this study were listed in Table S3.

Arabidopsis plants were grown on soil in a greenhouse under standard conditions. Seeds were stratified on water-imbibed paper in the dark at 4°C for 48 h, unless stated otherwise. 48 h-stratified seeds were germinated under continuous light at 25°C. When specified, germination was performed in the presence of 100 µM MG132 (Sigma-Aldrich), 1 µM ConA (Santa Cruz Biotechnology), 5 µM AZD-8055 (Santa Cruz Biotechnology), 100 µM cycloheximide (Sigma-Aldrich) or an equivalent concentration of DMSO (vehicle control).

### Total RNA isolation and reverse transcription quantitative PCR

Total RNA was isolated from seeds with phenol/chloroform 5:1 according to Di Berardino et al., 2018. Reverse transcription of isolated total RNA was realized using the M-MuLV Reverse Transcriptase kit (ThermoFisher Scientific) with an oligo (dT) primer according to the manufacturer’s protocol. The RT-qPCRs were run on a CFX 96 thermocycler (Biorad) using the primer pairs listed in Table S2. *ASAR1* (AT4G02080) and *MAP2B* (AT3G59990) were used as reference genes for the calculation of *NBR1* relative expression. All primers showed an efficiency of 100 ± 5%.

### Protein analysis using SDS-PAGE and immunoblotting

Total proteins were extracted from seeds or seedlings by homogenization in 2x NuPAGE LDS sample buffer (ThermoFischer Scientific) containing 0.1 M dithiotreitol (DTT) as described in Deruyffelaere et al., 2015. Protein extracts were heated at 37°C for 10 min prior to electrophoresis on Nu-PAGE gels using MOPS SDS running buffer according to the manufacturer’s instructions. Proteins were stained by Coomassie G250 according to Neuhoff et al. (1988). Coomassie-stained gels were

For immunoblot analysis, proteins were transferred onto polyvinylidene difluoride membranes (Millipore) and processed as described in D’Andrea et al. (2007). All primary antibodies were raised in rabbit. Antibodies directed against NBR1, actin, PEX14, VDAC1 and ARF1 were from Agrisera; ATG8A from Abcam; polyubiquitin chains linked at Lys-48 (UbK48 clone Apu2) and Lys-63 (UbK63 clone Apu3) from Millipore. Antibodies directed against OLE1 (anti rS3N) and OLE2 (anti rS4) were described in D’Andrea et al. (2007); against SLO1 (Anti AtHSD1) in d’Andréa et al. (2007); against CLO1 (Anti Clo1) in Purkrtova et al. (2007) and against LDAP1 in Coulon et al. (2024), respectively. To control protein loading, membranes were stained with amido-black 10B following immunodetection as described by Gershoni and Palade (1982). For immunoblot quantification, MultiGauge software (Fujifilm) was used.

### LD isolation and subcellular fractionation

For LD isolation, seeds or seedlings were ground in a seed homogenization buffer (SH buffer; 100 mM HEPES, pH 7.5, 500 mM sorbitol, 150 mM KCl, 1 mM MgCl_2_, 1 mM EDTA) supplemented with cOmplete™ mini EDTA-free protease inhibitors (Roche) using a glass-glass conical tissue grinder. LDs were isolated by ultracentrifugation on a discontinuous iodixanol density gradient and recovered from the floating fraction as described in Deruyffelaere et al. (2015).

For subcellular fractionation, a similar method as above was used with the following modifications: Seeds or seedlings were ground in SH-buffer lacking sorbitol (SO buffer) but containing 8% (w/v) sucrose and protease inhibitors. Organelles were separated using a continuous sucrose density gradient ranging from 0 to 48% (w/v) sucrose in SO buffer. Layers of 3 mL SO buffer containing 48%, 24% and 10% (w/v) sucrose were successively overlaid in an ultracentrifuge tube, which was then allowed to equilibrate on a glass bath for 3 h. The homogenate was carefully layered on top of the gradient and covered with 3 mL of SO buffer. After centrifugation at 100,000 *x g* for 18 h at 4 °C, the floating LD fraction and 0.8 mL of each remaining fraction were successively collected. Proteins were precipitated overnight by adding 180 µL of 100% (v/v) TCA and solubilized as described above.

### Fatty acid analysis by gas chromatography with flame ionization detection (GC-FID)

Fatty acid content and composition of seeds/seedlings were analyzed by GC-FID after direct transmethylation as described in Browse et al. (1986) with minor modifications described in Deruyffelaere et al. (2015).

### Accelerating aging

Accelerating aging was performed using the controlled deterioration protocol described in Rajjou et al. (2008). Briefly, seeds were equilibrated for one day in a closed tank (Van den Berg Klimaattechniek) at 30% relative humidity (RH) and 22 °C. Seeds were then subjected to 40 °C and 85% RH for 3, 5, 7 and 10 days before being dried back at 30 % RH and 22 °C. After drying, seeds were stored at 4 °C until use. Control seeds were equilibrated under the same conditions but immediately stored after equilibration. Germination assay was then carried out at 25 °C (16 h-light and 8 h-dark) for 8 days as described in Rajjou et al. (2008).

#### CLSM

Confocal observations were made with a SP5 II AOBS Tandem HyD confocal microscope (Leica) using a 63-water immersion objective. Neutral lipids were stained by incubating embryos with Bodipy 493/503 at 1.27 µg.mL^-1^ in water for 10 min, or HCS LipidTOX™ 637/655 Deep Red neutral lipid stain (Thermo Fischer Scientific) diluted 1:500 in water for 30 min, or Monodensylpentane (AUTODOT™ Visualization Dye (MDH) at 0.1 M in DMSO from CliniSciences) diluted 1:330 in water for 10 min. GFP, YFP, mCherry, Bodipy 493/503, HCS LipidTOX 637/655 and MDH were excited with the 488-nm line of an argon laser, 514 nm line of a diode laser, the 561-nm line of a diode laser, the 496-nm line of an argon laser, the 633-nm line of an helium/neon laser and the 405 nm line of a diode, respectively. Fluorescence emission was detected between 500 and 546 nm for GFP, 520 and 560 nm for YFP, 605 and 632 nm for mCherry, 500 and 547 nm for Bodipy 493/503, 637 and 652 nm for HCS LipidTOX 637/655 Deep Red and 420-480 nm for MDH. For multilabeling detections, a sequential line-scanning mode was used. The number of puncta carrying mCherry-NBR1 and/or GFP-ATG8f as well as the relative intensity profiles were determined using ImageJ Software (https:/rsbweb.nih.gov/ij/) developed by the National Institute of Health.

#### Pull-down assay

LDs were purified in triplicate from stratified seeds of Col-0 or p35S:mCherry-NBR1 lines, washed and resuspended in SH-buffer containing 0.05% (v/v) Nonidet^®^ P40 (Boehringer Mannheim) and cOmplete™ mini EDTA-free protease inhibitors. The OD_600_ of the LD fraction was used to adjust the volume of SH-buffer, ensuring that an equivalent amount of LDs was used in all samples. Pull down assay was performed using Magnetic RFP-Trap^®^_M (chromotek) according to the manufacturer’s instructions. Briefly, 60 µL of resuspended magnetic particles were incubated with 400 µL of the LD fraction for 2 h at 4 °C under gentle rotation. Supernatants were magnetically discarded and the beads were recovered and gently washed once with 500 µL of SH-buffer containing 0.01% (v/v) Nonidet^®^ P40 and protease inhibitors. Bound proteins were eluted by resuspending the beads with 2 x NuPAGE LDS sample buffer containing 0.1 M DTT and heating for 10 min at 75 °C. Eluted proteins were then processed for proteomic analysis.

#### Proteomic analysis

Proteins were loaded onto a polyacrylamide gel, and allowed to migrate for 1 cm by electrophoresis. Gels were cut into small pieces before trypsin in-gel digestion as previously described by Recorbet et al. (2021) and Langella et al. (2024) and provided in the Supplementary Methods. Briefly, following protein reduction and alkylation with iodoacetamide, gel pieces were digested overnight at 37°C using 100 ng of sequencing-grade modified trypsin (Promega). Peptides were subsequently extracted, dried under vacuum, and resuspended in 50 µL of 2% (v/v) acetonitrile (ACN) containing 0.1% (v/v) formic acid.

Liquid Chromatography–Mass Spectrometry (LC-MS)/MS analyses were performed using a timsTOF Pro mass spectrometer coupled to a nanoElute liquid chromatography system (Bruker Daltonik GmbH, Germany) and equipped with an Aurora C18 analytical column (IonOpticks, Australia). Data were acquired using the data-dependent acquisition parallel accumulation–serial fragmentation (dda-PASEF) method. Detailed chromatographic and mass spectrometry acquisition parameters are provided in the Supplementary Methods.

MS/MS data were searched against the *Arabidopsis thaliana* Araport11 protein database (48,360 entries, including the mCherry sequence) supplemented with a contaminant database using X!Tandem (version 2017.2.1.4; (Craig and Beavis, 2004)). Protein identification and inference were performed using i2MassChroQ (v1.0.11) (Langella et al., 2024)). Proteins were validated using a peptide E-value < 0.01, a protein log(E-value) < −4, and a minimum of two identified peptides per protein. For pull-down analyses, ubiquitination-related modifications and deamidation were additionally considered during database searches. Detailed database search parameters, including enzyme specificity, variable and fixed modifications, and mass tolerances, are provided in the Supplementary Methods.

Protein abundance was normalized by expressing the relative spectral intensity or protein abundance index (PAI) as a percentage of the total signal detected in each sample. Proteins showing more than a twofold enrichment relative to Col-0 and a Student’s *t*-test *p*-value ≤ 0.05 were considered significantly enriched. Statistical enrichment analysis of the proteomic datasets was performed using g:Profiler (g:GOST) (https://biit.cs.ut.ee/gprofiler/gost ; Kolberg et al., 2023).

Data post-processing and statistical analyses were performed using the R package MCQR (v0.6.15) (Balliau et al., 2025). Spectral count (SC) and extracted ion chromatogram (XIC) datasets were processed using dedicated bioinformatic workflows, including data filtering, normalization, missing value imputation, and statistical testing. Detailed processing parameters are provided in Supplementary Method. Differential protein abundance was assessed using adjusted *p*-value thresholds of <0.05 for SC analyses and <0.01 for XIC analyses. Descriptive analyses were performed using principal component analysis (PCA), heatmaps, and relative protein abundance ratios.

The bioinformatic tools used for proteomics analysis (X!TandemPipeline C++, MassChroQ, (Balliau et al., 2025)) are open and free resources available in the following repository: https://forgemia.inra.fr/pappso (Craig and Beavis, 2004). Proteomics analyses were performed on the PAPPSO platform (http://pappso.inra.fr) which is supported by INRAE, http://www.inrae.fr), the Ile-de-France regional council (https://www.iledefrance.fr/education-recherche), IBiSA (https://www.ibisa.net) and CNRS (http://www.cnrs.fr).

## Supporting information

Supplementary Data Set Table S1 A

Supplementary Data Set Table S1 B

Video S2 AZD ConA treatment

Video S2 AZD treatment

Video S2 ConA treatment

Video S2 DMSO treatment

## Acknowledgments

The authors would like to thank ChatGPT (OpenAI) and Gemini (Google) for assistance with the english translation and language refinement of this manuscript.

## Author contributions

I.B.M, C.D. S.D’A conceived, designed and performed experiments except AA experiment which was designed and conceived by L.R. and performed by B.C. I.B.M. and S.D’A wrote the manuscript. L. R. reviewed the manuscript. C.M.d’A reviewed the manuscript and provided materials. A. M. provided materials and performed experiments. M. F. reviewed the manuscript. All authors have read and agree to the final version of the manuscript.

## Data availability

The mass spectrometry proteomics data have been deposited to the ProteomeXchange Consortium (http://proteomecentral.proteomexchange.org) via the PRIDE partner repository (Perez-Riverol et al., 2024) with the dataset identifier PXD069683 for dataset Proteomic analysis and PXD069884 for dataset Pull-down analysis.

## Conflict of interest

The authors declare no conflict of interest.

## Funding

This work was partly supported by the French Agency for Research (Agence Nationale de la Recherche, grant no. ANR-21-CE20-0013 RecovOil Project).

## Supplementary Data

**Figure S1. AtNBR1 associates with LDs in the cotyledons of stratified and germinated seeds**

Cotyledons from stratified (0 h) and 24 h-germinated (24 h) seeds of Arabidopsis transgenic line expressing p35S:mCherry-NBR1 (mCh-NBR1, magenta) were imaged by CLSM after LD staining with Bodipy 493/503 (green). mCh-NBR1 localizes to LDs on two distinct structures, puncta (empty white arrowheads) and LD-surrounding rings (solid white arrowheads; magnified view in inset). Bars = 5 µm.

**Alt text: Figure S1:** AtNBR1 associates with LDs in the cotyledons of stratified and germinated seeds on two distinct structures : puncta and LD-surrounding rings.

**Figure S2. mCherry-NBR1 and GFP-ATG8f colocalize on puncta close to LDs**

The box plot with Tukey-style whiskers shows the number of puncta carrying mCherry-NBR1 (mCh-NBR1), GFP-ATG8f or both per image (n = 8 images). The red horizontal lines indicate the medians, while the bottom and top edges of the boxes represent the first (Q1) and third (Q3) quartiles, respectively. The percentage shows the proportion of puncta carrying both mCh-NBR1 and GFP-ATG8f among those carrying mCh-NBR1 (with or without GFP-ATG8f) or GFP-ATG8f (with or without mCh-NBR1).

**Alt text: Figure S2**: A comparative quantification of AtNBR1-positive puncta, GFP-ATG8f-positive puncta and both showed that mCherry-NBR1 and GFP-ATG8f colocalize on puncta close to LDs.

**Figure S3. Characterization of *nbr1* lines**

(**A**) Summary of Arabidopsis *nbr1* mutant lines, indicating the names used in previous publications and the nomenclature adopted in the present study. The previously unreported T-DNA mutant *nbr1-5* is characterized here. (**B**) Schematic representation of *NBR1* gene showing the position of the T-DNA insertion in *nbr1-5* mutant (position 12742418 on chromosome 4) and the position of the primers used for RT-PCR of *NBR1* mRNA. (**C**) *NBR1* mRNA in *nbr1-5* and Col-0 lines from 24-h-germinated seeds were detected by RT-PCR using NBR1-fwd4 and NBR1-rev3 (primers indicated by blue arrows on the gene structure). *EF1aA4* (*EF1α*) gene expression was used as a constitutive control, and *EF1α* mRNA were detected using primers EF-up and EF-down. The primer sequences are listed in Table S2. (**D**) AtNBR1 protein levels were analyzed in *nbr1* mutant lines by immunoblotting with anti-NBR1 antibodies. Total proteins extracted from equivalent amounts of stratified seeds from wild-type (Col-0) or mutant lines were loaded on each lane. The arrowhead indicates NBR1, and the asterisk marks a nonspecific band.

**Alt text**: **Figure S3**: Characterization of *nbr1* lines. **Figure S3A**: Summary of Arabidopsis *nbr1* mutant lines, indicating the names used in previous publications and the nomenclature adopted in the present study. **Figure S3B**: Schematic representation of *NBR1* gene showing the position of the T-DNA insertion in *nbr1-5* mutant and the position of the primers used for RT-PCR of *NBR1* mRNA. **Figure S3C**: *nbr1-5* line was depleted in *NBR1* transcripts. **Figure S3D**: *nbr1*-KO1 and *nbr1-5* lines were depleted in AtNBR1.

**Figure S4. Intravacuolar localization of NBR1-positive LD remnants**

Arabidopsis transgenic line co-expressing p35S:mCherry-NBR1 (magenta) and p35S:vac-gk (green) was analyzed by CLSM. Hypocotyls of 48 h-germinated seeds exposed to the drug (AZD + ConA) for 5 h were imaged after LD staining with Bodipy 493/503 (blue). Bars = 5 µm.

**Alt text: Figure S4:** NBR1-positive LD remnants were observed in hypocotyls of 48 h-germinated seeds treated with AZD + ConA for 5 h.

**Figure S5. Cycloheximide abolishes MG132-induced AtNBR1 accumulation**

Proteins were extracted from wild-type 42 h-germinated seeds treated for 17 h with DMSO, MG132, cycloheximide or both prior to recovery and analyzed by immunoblotting. AtNBR1 relative to actin content was measured using quantitative immunoblot (mean +/- SD from n = 3 samples of 50 seeds from independent plants). As cycloheximide inhibits germination, analyses were conducted separately for treatments lacking cycloheximide and those containing cycloheximide. Total proteins from an equivalent amount of seeds were loaded on each lane. Significant differences from the DMSO control were assessed using an exact one-tailed Mann–Whitney U test (p = 0.05). Individual values are represented by open circles, and significant differences are indicated by different letters.

**Alt text: Figure S5:** Treatment of germinated seeds with DMSO, MG132, Cycloheximide or MG132 + cycloheximide showed that cycloheximide abolishes MG132-induced AtNBR1 accumulation

**Figure S6. AtNBR1 degradation is delayed in *pux10-2* mutant**

Proteins were extracted from dry, stratified (0 h) or germinated seeds (24 to 72 h) of wild-type (Ws) or *pux10-2* mutant and analyzed by immunoblotting with anti-NBR1 antibodies. Total proteins from an equivalent amount of seeds were loaded on each lane.

**Alt text: Figure S6:** AtNBR1 accumulates in *pux10-2* during germination.

**Figure S7. AtNBR1 is not involved in a major LD degradation pathway**

(**A**) Fatty acid levels were comparable between *nbr1* mutants and Col-0 during germination. Fatty acids from dry, stratified (0 h) and germinated seeds (24 h and 48 h) of wild-type (Col-0), *nbr1*-KO1 and *nbr1-5* mutants were analyzed by GC-FID (mean +/- SD from n = 5 samples of 100 seeds from independent plants). (**B**) The absence of AtNBR1 has no effect on LD-integral protein degradation. Proteins were extracted from 48 h-germinated seeds (n = 2 independent plants) of wild-type (Col-0) or *nbr1* mutant lines (*nbr1*-KO1 and *nbr1-5),* analyzed by SDS-PAGE then immunoblotted with the specified antibodies. (**C**) The absence of AtNBR1 has no effect on OLE1 degradation. Proteins were extracted from 48 h-germinated seeds of wild-type (Col-0) or *nbr1* mutant lines (*nbr1*-KO1 and *nbr1-5),* analyzed by SDS-PAGE then immunoblotted with anti-OLE1 antibodies. Data are presented as individual biological replicates (n = 3 independent plants), with individual values represented by open circles. No statistically significant differences were detected using the Mann–Whitney U test (P > 0.05). Equal protein loading was verified by Coomassie G250 staining of a gel of the same protein extracts, as shown in the “Immunoblots used for quantification” section of the Supplementary Data.

**Alt text**: **Figure S7**: AtNBR1 is not involved in a major LD degradation pathway. **Figure S7A**: The absence of AtNBR1 in two independent *nbr1* mutants does not affect fatty acid levels, as determined by GC-FID analysis. **Figure S7B**: The absence of AtNBR1 does not affect LD-integral protein degradation in seeds from two independent *nbr1* mutants after 48 h of germination. **Figure S7C:** The absence of AtNBR1 does not affect OLE1 degradation in seeds from two independent *nbr1* mutants after 48 h of germination.

**Table S1. Putative AtNBR1 interactors identified by (A) LD pull-down from mCherry-NBR1 versus Col-0 and (B) comparative LD proteome of Col-0 versus *nbr1-KO1* mutant.**

**Alt text: Table S1:** By integrating two proteomic approaches, one single protein, TPP2, was identified as significantly enriched in LDs from Col-0 relative to *nbr1*-KO1 and in the pull-down of LDs from mCherry-NBR1 relative to Col-0.

**Table S2: Primers used in this study**

**Alt text: Table S2**: Primers used in this study.

**Table S3. Arabidopsis lines used in this study**

**Alt text: Table S3**: Arabidopsis lines used in this study

## Notes

http://proteomecentral.proteomexchange.org

### Competing Interest Statement

The authors have declared no competing interest.

