## Supplementary Data for "The selective autophagy receptor AtNBR1 associates with lipid droplets during seed germination and contributes to seed longevity"

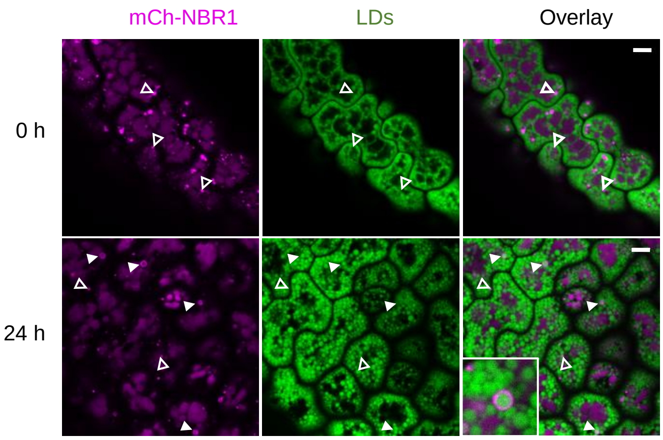


**Figure S1. AtNBR1 associates with LDs in the cotyledons of stratified and germinated seeds**

Cotyledons from stratified (0 h) and 24 h-germinated (24 h) seeds of Arabidopsis transgenic line expressing p35S:mCherry-NBR1 (mCh-NBR1, magenta) were imaged by CLSM after LD staining with Bodipy 493/503 (green). mCh-NBR1 localizes to LDs on two distinct structures, puncta (empty white arrowheads) and LD-surrounding rings (solid white arrowheads; magnified view in inset). Bars = 5 µm.
**Alt text: Figure S1:** AtNBR1 associates with LDs in the cotyledons of stratified and germinated seeds on two distinct structures : puncta and LD-surrounding rings.

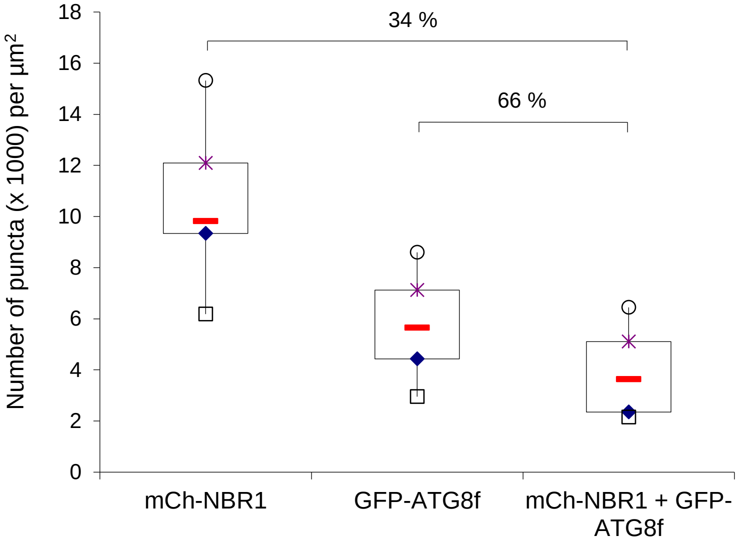


**Figure S2. mCherry-NBR1 and GFP-ATG8f colocalize on puncta close to LDs**

The box plot with Tukey-style whiskers shows the number of puncta carrying mCherry-NBR1 (mCh-NBR1), GFP-ATG8f or both per image (n = 8 images). The red horizontal lines indicate the medians, while the bottom and top edges of the boxes represent the first (Q1) and third (Q3) quartiles, respectively. The percentage shows the proportion of puncta carrying both mCh-NBR1 and GFP-ATG8f among those carrying mCh-NBR1 (with or without GFP-ATG8f) or GFP-ATG8f (with or without mCh-NBR1).
**Alt text: Figure S2**: A comparative quantification of AtNBR1-positive puncta, GFP-ATG8f-positive puncta and both showed that mCherry-NBR1 and GFP-ATG8f colocalize on puncta close to LDs.

**A**

| T-DNA mutant | Zhou et al. (2013) | Young et al. (2019) | Jung et al. (2020) | | This study |
|---|---|---|---|---|---|
| SALK_135513 | *nbr1-1* | *nbr1-1* | *nbr1-1* | - | - |
| GK_246H08 | *nbr1-2* | *nbr1-2* | *nbr1-2* | - | *nbr1-2* |
| SALK_144852 | - | *nbr1-1* | *nbr1-3* | - | *nbr1-3* |
| WiscDsLoxHs007_07A | - | *nbr1-4* | *nbr1-4* | - | *nbr1-4* |
| GK_281A10 | - | - | - | - | *nbr1-5* |
| **CRISPR/Cas 9** | | | | **Tarnowski et al. (2020a)** | |
| KO1 : 1062 nt Δ (from -133 to +929) | - | - | - | KO1 | *nbr1*-KO1 |

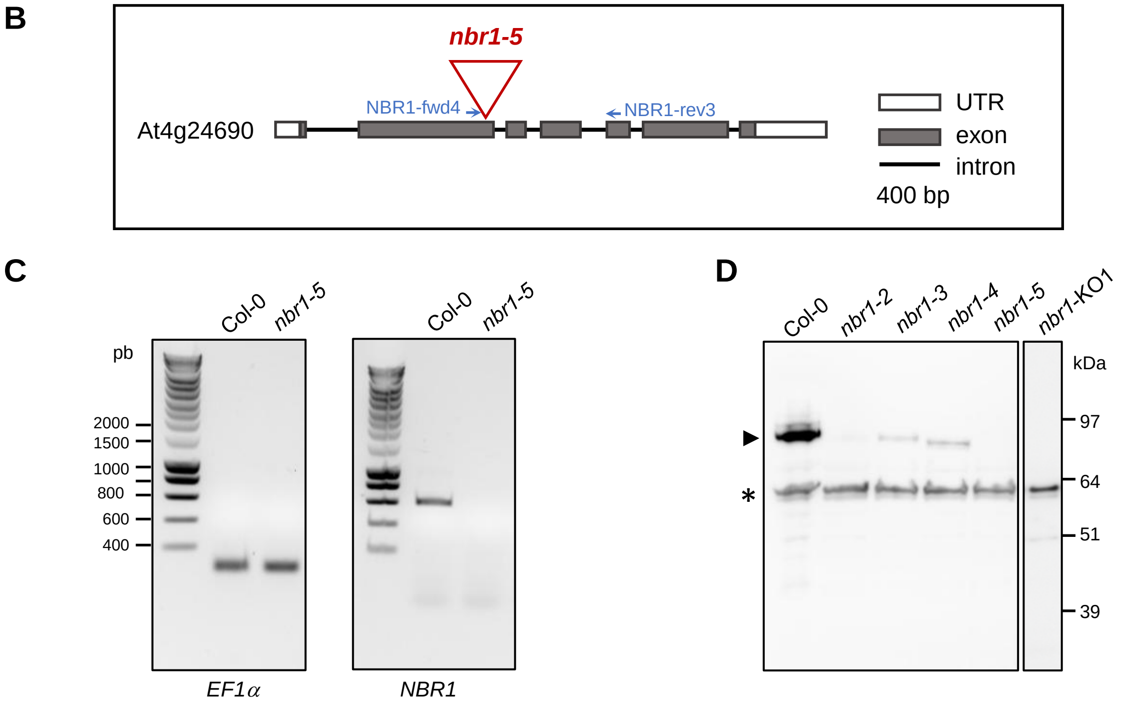


**Figure S3. Characterization of *nbr1* lines**

(**A**) Summary of Arabidopsis *nbr1* mutant lines, indicating the names used in previous publications and the nomenclature adopted in the present study. The previously unreported T-DNA mutant *nbr1-5* is characterized here. (**B**) Schematic representation of *NBR1* gene showing the position of the T-DNA insertion in *nbr1-5* mutant (position 12742418 on chromosome 4) and the position of the primers used for RT-PCR of *NBR1* mRNA. (**C**) *NBR1* mRNA in *nbr1-5* and Col-0 lines from 24-h-germinated seeds were detected by RT-PCR using NBR1-fwd4 and NBR1-rev3 (primers indicated by blue arrows on the gene structure). *EF1aA4* (*EF1α*) gene expression was used as a constitutive control, and *EF1α* mRNA were detected using primers EF-up and EF-down. The primer sequences are listed in Table S2. (**D**) AtNBR1 protein levels were analyzed in *nbr1* mutant lines by immunoblotting with anti-NBR1 antibodies. Total proteins extracted from equivalent amounts of stratified seeds from wild-type (Col-0) or mutant lines were loaded on each lane. The arrowhead indicates NBR1, and the asterisk marks a nonspecific band.

**Alt text**: **Figure S3**: Characterization of *nbr1* lines. **Figure S3A**: Summary of Arabidopsis *nbr1* mutant lines, indicating the names used in previous publications and the nomenclature adopted in the present study. **Figure S3B**: Schematic representation of *NBR1* gene showing the position of the T-DNA insertion in *nbr1-5* mutant and the position of the primers used for RT-PCR of *NBR1* mRNA. **Figure S3C**: *nbr1-5* line was depleted in *NBR1* transcripts. **Figure S3D**: *nbr1*-KO1 and *nbr1-5* lines were depleted in AtNBR1.

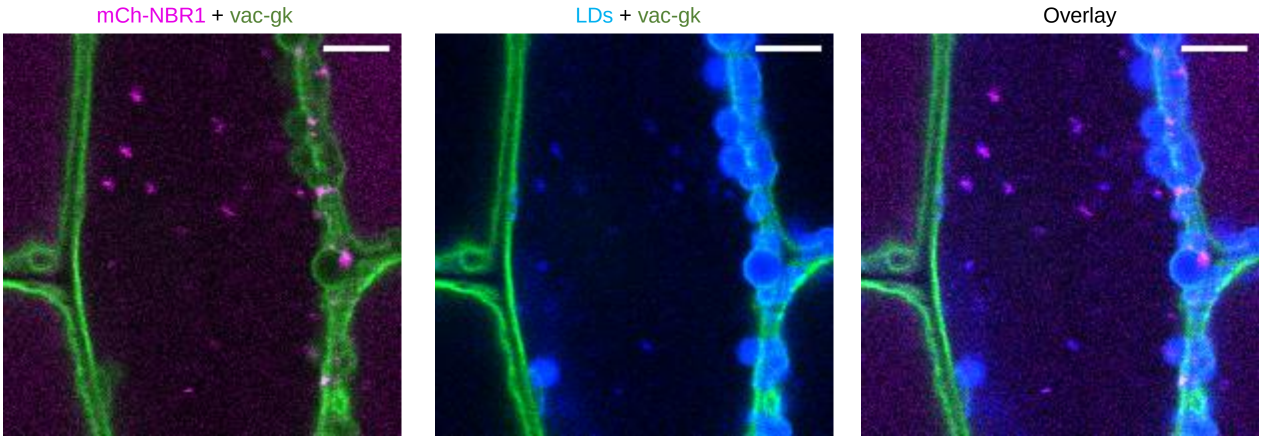


**Figure S4. Intravacuolar localization of NBR1-positive LD remnants**

Arabidopsis transgenic line co-expressing p35S:mCherry-NBR1 (magenta) and p35S:vac-gk (green) was analyzed by CLSM. Hypocotyls of 48 h-germinated seeds exposed to the drug (AZD + ConA) for 5 h were imaged after LD staining with Bodipy 493/503 (blue). Bars = 5 µm.
**Alt text: Figure S4:** NBR1-positive LD remnants were observed in hypocotyls of 48 h-germinated seeds treated with AZD + ConA for 5 h.

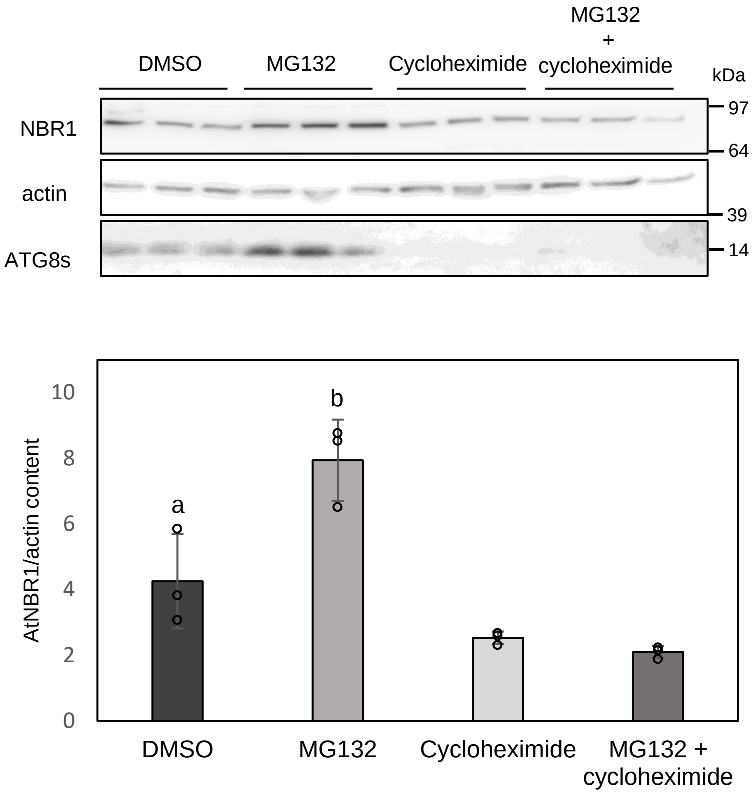


**Figure S5**. **Cycloheximide abolishes MG132-induced AtNBR1 accumulation**

Proteins were extracted from wild-type 42 h-germinated seeds treated for 17 h with DMSO, MG132, cycloheximide or both prior to recovery and analyzed by immunoblotting. AtNBR1 relative to actin content was measured using quantitative immunoblot (mean +/- SD from n = 3 samples of 50 seeds from independent plants). As cycloheximide inhibits germination, analyses were conducted separately for treatments lacking cycloheximide and those containing cycloheximide. Total proteins from an equivalent amount of seeds were loaded on each lane. Significant differences from the DMSO control were assessed using an exact one-tailed Mann–Whitney U test (p = 0.05). Individual values are represented by open circles, and significant differences are indicated by different letters.
**Alt text: Figure S5:** Treatment of germinated seeds with DMSO, MG132, Cycloheximide or MG132 + cycloheximide showed that cycloheximide abolishes MG132-induced AtNBR1 accumulation.

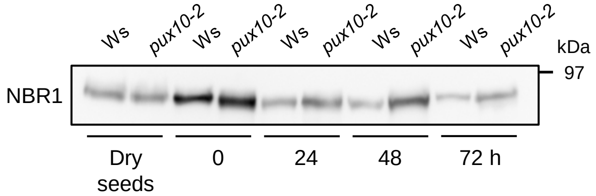


**Figure S6**. **AtNBR1 degradation is delayed in *pux10-2* mutant**

Proteins were extracted from dry, stratified (0 h) or germinated seeds (24 to 72 h) of wild-type (Ws) or *pux10-2* mutant and analyzed by immunoblotting with anti-NBR1 antibodies. Total proteins from an equivalent amount of seeds were loaded on each lane.
**Alt text: Figure S6:** AtNBR1 accumulates in *pux10-2* during germination.

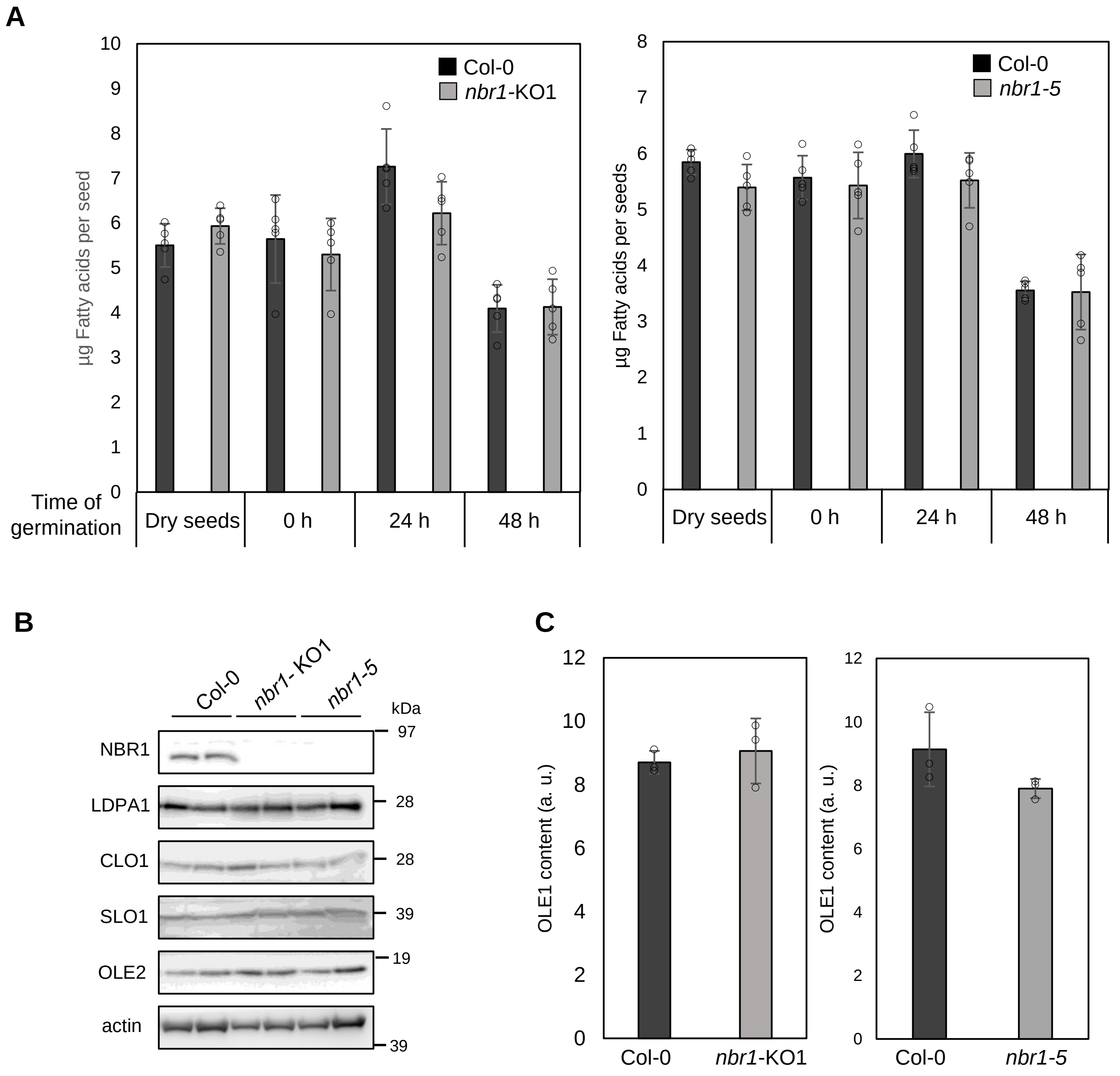


**Figure S7. AtNBR1 is not involved in a major LD degradation pathway**

(**A**) Fatty acid levels were comparable between *nbr1* mutants and Col-0 during germination. Fatty acids from dry, stratified (0 h) and germinated seeds (24 h and 48 h) of wild-type (Col-0), *nbr1*-KO1 and *nbr1-5* mutants were analyzed by GC-FID (mean +/- SD from n = 5 samples of 100 seeds from independent plants). (**B**) The absence of AtNBR1 has no effect on LD-integral protein degradation. Proteins were extracted from 48 h-germinated seeds (n = 2 independent plants) of wild-type (Col-0) or *nbr1* mutant lines (*nbr1*-KO1 and *nbr1-5),* analyzed by SDS-PAGE then immunoblotted with the specified antibodies. (**C**) The absence of AtNBR1 has no effect on OLE1 degradation. Proteins were extracted from 48 h-germinated seeds of wild-type (Col-0) or *nbr1* mutant lines (*nbr1*-KO1 and *nbr1-5),* analyzed by SDS-PAGE then immunoblotted with anti-OLE1 antibodies. Data are presented as individual biological replicates (n = 3 independent plants), with individual values represented by open circles. No statistically significant differences were detected using the Mann–Whitney U test (P > 0.05). Equal protein loading was verified by Coomassie G250 staining of a gel of the same protein extracts, as shown in the "Immunoblots used for quantification" section of the Supplementary Data.

**Alt text**: **Figure S7**: AtNBR1 is not involved in a major LD degradation pathway. **Figure S7A**: The absence of AtNBR1 in two independent *nbr1* mutants does not affect fatty acid levels, as determined by GC-FID analysis. **Figure S7B**: The absence of AtNBR1 does not affect LD-integral protein degradation in seeds from two independent *nbr1* mutants after 48 h of germination. **Figure S7C:** The absence of AtNBR1 does not affect OLE1 degradation in seeds from two independent *nbr1* mutants after 48 h of germination.

**Table S1. Putative AtNBR1 interactors identified by (A) LD pull-down from mCherry-NBR1 versus Col-0 and (B) comparative LD proteome of Col-0 versus *nbr1-KO1* mutant.**

| Function [(a)] | GENE ID | Protein name | Description [(a)] | (A) LD Pull-down mCh-NBR1 vs. Col-0 [(b)] | (B) LD proteome Col-0 vs. *nbr1*-KO1 [(c)] |
|---|---|---|---|---|---|
| **Protein biosynthesis** | | | | | |
| | AT1G58380 | RPS2A | Small ribosomal subunit protein uS5z | 6.4 | - |
| | AT4G34670 | RPS3AB | Small ribosomal protein eS1y | 4.9 | - |
| | AT3G04840 | RPS3AA | Small ribosomal subunit protein eS1z | 3.5 | - |
| | AT1G23410 | R27AA | Ubiquitin-ribosomal protein eS31z fusion protein | 2.3 | - |
| | AT2G42740 | RPL11A | Large ribosomal subunit protein uL5z | - | 2.5 |
| | AT1G50200 | ALATS | Alanine--tRNA ligase | - | not in *nbr1*-KO1 |
| | AT3G13460 | ECT2 | YTH domain-containing protein | - | not in *nbr1*-KO1 |
| **Protein folding** | | | | | |
| | AT1G59860 | HSP17.6A | 17.6 kDa class I heat shock protein 1 | 11.6 | - |
| | AT1G53540 | HSP17.6C | 17.6 kDa class I heat shock protein 3 | 2.2 | - |
| | AT3G46230 | HSP17.4A | 17.4 kDa class I heat shock protein | 2.2 | - |
| | AT2G29500 | HSP17B | 17.6 kDa class I heat shock protein 2 | 2.0 | - |
| | AT2G16595 | TRAP2C | Translocon-associated protein (TRAP)2C alpha subunit | 5.9 | - |
| | AT5G22060 | ATJ2 | Chaperone protein dnaJ 2 | - | 9.7 |
| | AT2G35720 | ATJ13 | Chaperone protein dnaJ 13 | - | not in *nbr1*-KO1 |
| | AT3G03960 | CCT8 | T-complex protein 1 subunit theta | - | 2.4 |
| | AT1G56340 | CRT1 | Calreticulin 1a | - | not in *nbr1*-KO1 |
| **Protein degradation** | | | | | |
| | **AT4G20850** | **TPP2** | **Tripeptidyl peptidase 2** | **2.5** | **2.7** |
| | AT5G38470 | RAD23D | Ubiquitin receptor RAD23d | - | not in *nbr1*-KO1 |
| **Membrane organization and dynamics** | | | | | |
| | AT1G07985 | LIPA | LD-plasma membrane adaptor | 8.8 | - |
| | AT2G28900 | OEP161 | Outer envelope pore protein 16-1 | 2.8 | - |
| | AT5G42570 | B-cell receptor-associated 31-like protein | Endoplasmic reticulum transmembrane protein | 2.5 | - |
| | AT2G47770 | TSPO | Translocator protein homolog | 2.1 | - |
| | AT2G01470 | STLP2 | SEC12-like protein 2 | 2.1 | - |
| | AT5G41600 | RTNLB4 | Reticulon-like protein B4 | - | 3.8 |
| | AT1G22610 | MCTP6 | Multiple C2 domain and transmembrane region protein 6 | - | not in *nbr1*-KO1 |
| **Metabolic enzymes** | | | | | |
| | AT5G65750 | MPA24.10 | 2-oxoglutarate dehydrogenase | - | 3.7 |
| | AT3G48000 | ALDH2B4 | Aldehyde dehydrogenase 2B4 | - | 3.5 |
| | AT1G64190 | PGD1 | 6-phosphogluconate dehydrogenase family protein | - | 2.9 |
| | AT2G30200 | MCAT | Malonyl-CoA:ACP malonyltransacylase | - | 2.7 |
| | AT1G53240 | MDH1 | Malate dehydrogenase 1 | - | 2.6 |
| | AT3G17810 | PYD1 | Dihydropyrimidine dehydrogenase | - | not in *nbr1*-KO1 |
| | AT3G25800 | PP2AA2 | Protein phosphatase 2A subunit A2 | - | not in *nbr1*-KO1 |
| | AT3G02230 | RGP1 | UDP-arabinopyranose mutase 1 | - | not in *nbr1*-KO1 |
| | AT3G08510 | PLC2 | Phospholipase C 2 | - | not in *nbr1*-KO1 |
| | AT5G18170 | GDH1 | Glutamate dehydrogenase 1 | - | not in *nbr1*-KO1 |
| | AT5G08530 | NADH | NADH dehydrogenase [ubiquinone] flavoprotein 1 | - | not in *nbr1*-KO1 |
| **Other or unknown functions** | | | | | |
| | AT1G09570 | PHYA | Phytochrome A | 21.4 | - |
| | AT4G32470 | QCR7-1 | QCR7-1 Cytochrome b-c1 complex subunit 7-1 | 4.5 | - |
| | AT4G00090 | Transducin/WD40 repeat-like superfamily protein | Transducin/WD40 repeat-like superfamily protein | 2.7 | - |
| | AT1G33970 | IAN9 | Immune-associated nucleotide-binding protein 9 | not in Col-0 | - |
| | AT5G54540 | MRB17-4 | MRB17-4 | not in Col-0 | - |
| | AT5G40770 | PHB3 | Prohibitin 3 | - | 8.7 |
| | AT4G28510 | PHB1 | Prohibitin 1 | - | not in *nbr1*-KO1 |
| | AT1G54780 | Thylakoid lumen 18.3 kDa protein | Thylakoid lumen 18.3 kDa protein | - | 5.2 |
| | AT1G32400 | TOM2A | Tobamovirus multiplication protein 2A | - | 2.5 |
| | AT3G14650 | CYP72A11 | Cytochrome P450 72A11 | - | not in *nbr1*-KO1 |
| | AT2G02470 | ALFINL6 | PHD finger protein alfin-like 6 | - | not in nbr1-KO1 |

[(a)] based on UniProt annotations (www.uniprot.org) ; [(b)] Significant enrichment ($p<0.05$) or not detected in Col-0; [(c)] Significant enrichment ($p<0.05$) or not detected in *nbr1*-KO1.

**Alt text: Table S1**: By integrating two proteomic approaches, one single protein, TPP2, was identified as significantly enriched in LDs from Col-0 relative to *nbr1*-KO1 and in the pull-down of LDs from mCherry-NBR1 relative to Col-0.

**Table S2 : Primers used in this study**

| Type | Primer name | Primer sequence |
|---|---|---|
| RT-PCR | NBR1-fwd4 | GCATCCTCGTCGTGTTTGTCAT |
| RT-PCR | NBR1-rev3 | CCAGGTAACTCTGGTGCAACA |
| RT-PCR | EF-up | CTGGAGGTTTTGAGGCTGGTAT |
| RT-PCR | EF-down | CCAAGGGTGAAAGCAAGAAGA |
| RT-qPCR | NBR1-rev | ACACGACGAGGATGCTTGTT |
| RT-qPCR | NBR1-for | CCACTGGCGCTCATTCAAAG |
| RT-qPCR | MAP2B-for | GTGACTCTGGCATTGTTCAG |
| RT-qPCR | MAP2B-rev | AAACTGGATACATAGCTTCCTTTC |
| RT-qPCR | ASAR1-rev | AAAGCTAGGTACGGTTTAAGAC |
| RT-qPCR | ASAR1-for | GCTGTGTTATTATTAAGCCGTTAAG |

**Alt text: Table S2**: Primers used in this study.

#### Table S3. Arabidopsis lines used in this study

| Plant lines | Description | Reference |
|---|---|---|
| ***pux10-1*** | TDNA SAIL_1187 | Deruyffelaere *et al.*, 2018 |
| ***pux10-2*** | TDNA FLAG_211E02 | Deruyffelaere *et al.*, 2018 |
| ***atg2-1*** | TDNA SALK_076727 | Yoshimoto *et al.*, 2009 |
| ***atg4a4b-1*** | TDNA | Yoshimoto *et al.*, 2004 |
| ***nbr1-2*** | TDNA GK_246H08 | Zhou *et al.,* 2013 |
| ***nbr1-3*** | TDNA SALK_144852 | Jung *et al.*, 2020 |
| ***nbr1-4*** | TDNA WiscDsLoxHs007_07A | Young *et al.,* 2019 |
| ***nbr1-5*** | GK_281A10 | This study |
| ***nbr1*-KO1** | CRISPER/Cas9 | Tarnowski *et al.*, 2020 |
| **mCherry-NBR1** | p35S:mCherry-NBR1 | Svenning *et al.*, 2011 |
| **GFP-ATG8f** | pactin:GFP-ATG8f | Di Berardino *et al.,* 2018 |
| **PUX10-GFP** | pPUX10:PUX10-GFP | Deruyffelaere *et al.*, 2018 |
| **YFP-CDC48A** | pCDC48:YFP-CDC48A | Deruyffelaere *et al.*, 2018 |
| **vac-gk** | p35S:vac-gk | Nelson *et al.,* 2007 |
| **mCherry-NBR1/PUX10-GFP** | p35S:mCherry-NBR1 / pPUX10:PUX10-GFP | This study |
| **mCherry-NBR1/YFP-CDC48A** | p35S:mCherry-NBR1 / pCDC48A:YFP-CDC48A | This study |
| **mCherry-NBR1/GFP-ATG8f** | p35S:mCherry-NBR1 / pactin:GFP-ATG8f | This study |
| **mCherry-NBR1/vac-gk** | p35S:mCherry-NBR1 / p35S:vac-gk | This study |

**Alt text: Table S3**: Arabidopsis lines used in this study.

### Immunoblots used for quantification

#### Figure 1B

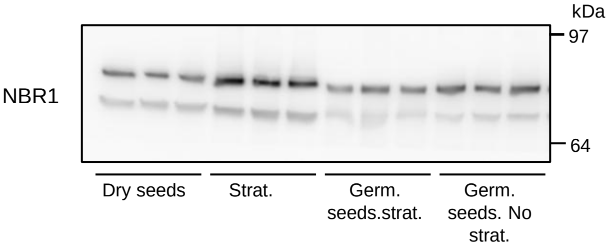


#### Figure 4B

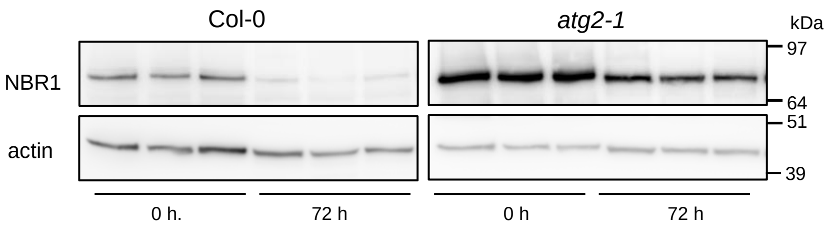


#### Figure S7C

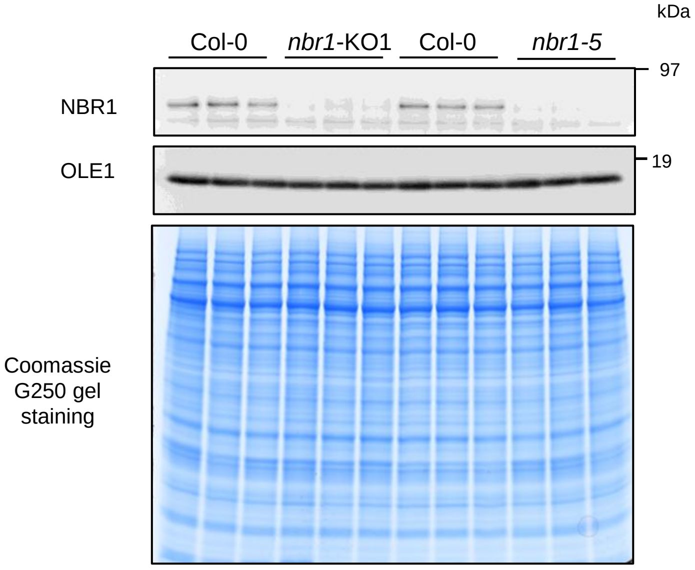
