## Supplementary methods for "The selective autophagy receptor AtNBR1 associates with lipid droplets during seed germination and contributes to seed longevity"

**LC MS/MS analysis**

**In gel digestion**

Gel pieces were first destained by incubation in 50 µL of 40% (v/v) ethanol and 10% (v/v) acetic acid for 30 min, followed by incubation in 50 µL of 50 mM NH₄HCO₃ containing 50% (v/v) acetonitrile (ACN) for 30 min. This latter step was repeated three times. Gel pieces were then dehydrated in 100% (v/v) ACN for 15 min. Proteins were reduced by incubation in 10 mM dithiothreitol (DTT) prepared in 50 mM NH₄HCO₃ for 30 min at 56°C, and subsequently alkylated with 55 mM iodoacetamide in 50 mM NH₄HCO₃ for 45 min at room temperature in the dark. Gel pieces were rinsed with 50 µL of 50 mM NH₄HCO₃ containing 50% (v/v) ACN for 30 min and dehydrated again in 100% (v/v) ACN for 15 min. In-gel digestion was performed overnight at 37°C using 100 ng of trypsin diluted in 50 mM NH₄HCO₃, with sufficient volume added to completely cover the gel pieces. The following day, the digestion supernatant was transferred to a fresh tube. Peptides were extracted from the gel by incubation in 50 µL of 0.5% (v/v) trifluoroacetic acid (TFA) and 50% ACN (v/v) for 30 min, and the extract was pooled with the digestion supernatant. A second extraction was then performed using 100% ACN (v/v). The combined peptide extracts were dried in a SpeedVac concentrator and resuspended in 50 µL of loading buffer containing 2% (v/v) ACN and 0.1% (v/v) formic acid prior to LC–MS/MS analysis.

**LC/MS-MS acquisition**

Samples were loaded onto an Acclaim™ PepMap™ C18 trap column (0.1 mm i.d. × 20 mm, 5 µm particle size, 100 Å pore size; Thermo Fisher Scientific) and washed with loading buffer. The trap column was then connected to an Aurora C18 analytical column (75 µm i.d. × 250 mm, 1.6 µm particle size, 120 Å pore size; IonOpticks, Australia) for peptide separation. For proteomic experiments, peptides were eluted at a flow rate of 200 nL min⁻¹ using the following gradient of buffer B (0.1% (v/v) formic acid in ACN) : 2–5% buffer B over 1 min, 5–13% over 18 min, 13–19% over 7 min, and 19–22% over 4 min. For pull-down experiments, peptides were eluted at a flow rate of 200 nL min⁻¹ using the following gradient: 2–12% buffer B over 18 min, 12–20% over 7 min, and 20–30% over 4 min. For both, the total run time was 40 min, including a column regeneration step at 95% buffer B for 7 min. The column temperature was maintained at 50°C. Peptides were introduced into the mass spectrometer using a CaptiveSpray nano-electrospray ionization source (Bruker Daltonik GmbH) operated at an electrospray voltage of 1.6 kV, a source temperature of 180°C, and a dry gas flow rate of 3 L min⁻¹. Data were acquired in data-dependent acquisition parallel accumulation–serial fragmentation (dda-PASEF) mode. A total of six PASEF scans were performed per acquisition cycle, targeting precursor ions with charge states ranging from 0 to 5 and resulting in a total cycle time of 1.30 s. Each PASEF scan was acquired with a ramp time of 180 ms. Ion mobility was calibrated using Agilent ESI-L Tune Mix over a mobility range of 0.6–1.3 V·s cm⁻² (1/K₀). The ion mobility scan range used for data acquisition was 0.7–1.10 V·s cm⁻² (1/K₀). Collision energy was applied as a linear function of ion mobility, increasing from 20.0 eV at 0.60 V·s cm⁻² (1/K₀) to 59.0 eV at 1.60 V·s cm⁻² (1/K₀). MS and MS/MS spectra were acquired over an m/z range of 100–1700. The TIMS accumulation time was set to 180 ms. Precursor selection was performed using a target intensity of 14,000 arbitrary units (a.u.) and a minimum intensity threshold of 1,000 a.u. Absolute intensity thresholds were set to 10 counts for mass spectrum peak detection and 5,000 counts for mobilogram peak detection.

**Protein identification**

Database searches were performed using the X!tandem (Craig 2024 ; version 2017.2.1.4) against the *arabidopsis thaliana* database with 48,360 proteins entries (Araport version 11 release 201606 with mCherry Sequence) and a contaminant database. Enzymatic cleavage was set to trypsin with proline rules, allowing one missed cleavage in the first-pass search. Carbamidomethylation of cysteine residues was set as static modification. Oxidation of methionine, protein N-terminal acetylation (with or without initiator methionine excision), and pyroglutamate formation from N-terminal glutamine or glutamic acid residues were included as variable modifications. For the pull-down analysis, ubiquitination-related modifications on lysine residues (tetrapeptidyl-lysine and dipeptidyl-lysine) as well as deamidation of glutamine and asparagine residues were included as variable modifications during the refinement step. Precursor and fragment mass tolerance were 20 ppm. Identified proteins were filtered and infered using i2MassChroQ (version 1.0.11, <http://pappso.inrae.fr/>) (Langella et al., 2024) Data filtering was achieved according to a peptide E-value <0.01, protein log (E-value) < –4 and to a minimum of two identified peptides per protein.

**Statistics analysis**

Data post-processing and statistical analyses were performed using the R package MCQR (v0.6.15) (Balliau et al., 2025). Distinct bioinformatic workflows were applied to spectral count (SC) and extracted ion chromatogram (XIC) datasets. For SC-based analyses, the workflow included: (i) removal of proteins represented by fewer than five spectra across all samples, and (ii) removal of proteins displaying less than a 1.5-fold abundance variation between strains. For XIC-based analyses, the workflow included: (i) removal of peptides exhibiting high retention time variability (>30 s) or peak widths exceeding 300 s; (ii) normalization of peptide intensities using a reference sample; (iii) exclusion of shared peptides; (iv) removal of peptides with more than 10% missing values across the entire dataset; (v) exclusion of proteins identified by fewer than two unique peptides; (vi) imputation of missing peptide intensity values by replacing them with the minimum abundance measured for the corresponding protein across the dataset; and (vii) removal of peptides exhibiting less than a 1.5-fold abundance variation between strains. Protein abundance changes were considered statistically significant when the adjusted p-value was <0.05 for SC analyses and <0.01 for XIC analyses.
