## Supplementary Video Legends for "The selective autophagy receptor AtNBR1 associates with lipid droplets during seed germination and contributes to seed longevity"

**Video S1. Mobile NBR1-positive LD aggregates within the vacuole.**

Arabidopsis transgenic line expressing p35S:mCherry-NBR1 (magenta) was analyzed by CLSM. Hypocotyls of 48 h-germinated seeds were imaged after LD staining with Bodipy 493/503 (green). Forty frames were recorded for around 2 minutes and saved under 7 frames per sec. Bars = 5 µm.

**Video S2. Effect of autophagy modulation on mobile NBR1-positive LDs within the vacuole.**

Arabidopsis transgenic line expressing p35S:mCherry-NBR1 (magenta) was analyzed by CLSM. Hypocotyls of 48 h-germinated seeds exposed to the drugs (ConA, AZD or AZD+ConA) or DMSO for 5 h were imaged after LD staining with Bodipy 493/503 or MDH (green). Forty frames were recorded for around 2 minutes and saved under 7 frames per sec. Bars = 5 µm.

**Video S3. Intravacuolar localization of mobile NBR1-positive LD remnants.**

Arabidopsis transgenic line co-expressing p35S:mCherry-NBR1 (magenta) and p35S:vac-gk (green) was analyzed by CLSM. Hypocotyls of 48 h-germinated seeds exposed to the drug (AZD + ConA) for 5 h were imaged after LD staining with Bodipy 493/503 (blue). Forty frames were recorded for around 2 minutes and saved under 7 frames per sec. Bars = 5 µm.
